# CRISPR-Cas effector specificity and target mismatches determine phage escape outcomes

**DOI:** 10.1101/2022.05.30.494023

**Authors:** Michael A. Schelling, Giang T. Nguyen, Dipali G. Sashital

## Abstract

CRISPR-mediated interference relies on complementarity between a guiding CRISPR RNA (crRNA) and target nucleic acids to provide defense against bacteriophage. Phages escape CRISPR-based immunity mainly through mutations in the PAM and seed regions. However, previous specificity studies of Cas effectors, including the class 2 endonuclease Cas12a, have revealed a high degree of tolerance of single mismatches. The effect of this mismatch tolerance has not been extensively studied in the context of phage defense. Here, we tested defense against lambda phage provided by Cas12a-crRNAs containing pre-existing mismatches against the genomic targets in phage DNA. We observe a correlation between Cas12a mismatch tolerance in vitro and phage defense on solid media. However, in liquid media, we find that most pre-existing crRNA mismatches lead to phage escape and lysis, regardless of whether the mismatches ablate Cas12a cleavage in vitro. We used high-throughput sequencing to examine the target regions of phage genomes following CRISPR challenge. Mismatches at all locations in the target accelerated emergence of mutant phage, including mismatches that greatly slowed cleavage in vitro. Mutations arose near the existing mismatch, in some cases resulting in multiple PAM-distal mismatches allowing for phage escape. Similar experiments with Cas9 showed the location of emergent target mutations was unaffected by pre-existing crRNA-target mismatches. Expression of multiple mismatched crRNAs prevented new mutations from arising in multiple targeted locations, allowing Cas12a mismatch tolerance to provide stronger and longer term protection. These results demonstrate that Cas effector mismatch tolerance and existing target mismatches strongly influence phage evolution.

## Introduction

CRISPR-Cas (clustered regularly interspaced short palindromic repeats-CRISPR associated) systems are adaptive immune systems found in bacteria and archaea^1–3^. These systems use ribonucleoprotein effector complexes to find and destroy foreign nucleic acids that have entered the cell. CRISPR effector complexes are guided by a CRISPR RNA (crRNA) to a nucleic acid target that is complementary to a section of the crRNA called the spacer. Bacteria can acquire new spacer sequences that allow them to mount an immune response against threats they have not previously encountered^4, 5^.

An important function of CRISPR-Cas systems is to prevent infection by bacteriophages, which can have significant impact on the composition of a bacterial population^6–9^. As a CRISPR effector complex requires a match between its crRNA and a target to engage in interference, selection occurs for phages with mutations in targeted genomic regions^10–12^. Mutations in CRISPR targets create mismatches between the target and the crRNA that weaken the base-pairing interaction^13–15^, slowing or stopping target matching by Cas effectors^16^ and allowing phages to safely multiply in the bacterial cell. Different CRISPR-Cas systems have DNA or RNA as a primary target and prevent infection at the cellular and population level^17–22^. Target binding is more stringent in DNA targeting systems, mitigating highly damaging off-target cleavage of host DNA^23^. In these systems, a protospacer adjacent motif (PAM) next to the target is required to initiate base pairing^24–27^. Complete base pairing is especially important in the region next to the PAM, called the seed region^28–34^. Accordingly, mutations that allow phages to escape CRISPR immunity are often single mutations in the PAM or seed region^10–12, 35^.

There have been multiple proposed but non-competing mechanisms for this mutagenesis. Mutants may exist due to natural genetic variation in the population and these could be selected through CRISPR pressure and become dominant in the population over time^11, 36^. Alternatively, escape mutations may be generated by Cas effector cleavage and subsequent error prone DNA repair^37^. It has been shown that cleavage by Cas effectors causes large deletions to appear in the genome of T4 phage, resulting in loss of the crRNA target sequence^38^. Lambda phage encoded DNA repair enzymes have been implicated in generating mutations in CRISPR targets that allow escape^39^. It remains unclear to what degree each of these mutagenesis pathways contribute to phage escape under different conditions.

Escape mutations are evident in natural settings as bacterial CRISPRs often contain mismatched spacers to common mobile genetic elements and the genomes of phages^27, 40–43^ . New spacers are added at the leader end of CRISPR arrays and these new spacers are more likely to match a target in a phage genome exactly^44, 45^. Genomic evidence also shows that spacer sequences in a CRISPR array do not commonly develop mutations, and are fixed once they are acquired^12, 46^. Instead, spacers are lost from the array entirely when they lose effectiveness as mutations accumulate in targeted genomic elements.

Mismatched spacers may provide some benefit to the host. Spacers against mutated targets drive some Cas effectors towards primed spacer acquisition, in which new spacers are preferentially acquired from genomes targeted by the Cas effector^11, 31, 47–50^. Mismatched crRNAs may also provide low level immunity through continued target cleavage. Cas effectors tolerate mismatches between the crRNA and target, allowing cleavage of mutated targets^26, 28, 29, 51–55^. This lax specificity may partially prevent phage escape. The type V-A Cas12a effector has been shown to tolerate multiple mismatches between its guiding crRNA and the target in vitro^53, 56, 57^ leading to rare off-target genome edits in cells^57–61^. However, this mismatch tolerance varies depending on the crRNA sequence and type of mismatch. Multiple Cas12a variants also have the ability to nick double stranded DNA targets with many (3-4) mismatches^53, 62^. These in vitro observations raise the question of how the specificity of Cas12a affects its role in preventing infection by phage with target mutations.

To test this, we subjected bacteria expressing Cas12a and crRNAs with varying target mismatches to phage infection. We found unexpected discrepancies between the effect of crRNA mismatches on target cleavage in vitro and survival of bacteria in liquid culture. This led us to monitor mutant emergence in phage populations. Using high-throughput sequencing, we discovered enrichment of a large variety of mutations when the phage was targeted by different crRNAs with and without mismatches. Single crRNA mismatches, even those outside of the seed region, had a drastic effect on the ability of bacteria to survive phage exposure, demonstrating the importance of spacer diversity as mutations in target genomic regions propagate. crRNA mismatches increased the rate at which mutant phage arose in the population. Mutation types that arose depended not only on the location of pre-existing mismatches but also on the location of the target in the genome. The diversity of escape mutations we observed was unique to Cas12a as crRNA mismatches did not affect the range of mutations that allowed escape from Cas9 targeting. We found that phage populations evolve in different ways to resist CRISPR interference depending on Cas effector specificity, existing crRNA-target mismatches, and the location of CRISPR targets in the phage genome.

## Results

### crRNA mismatches throughout the spacer decrease phage protection provided by Cas12a

To investigate the effect of crRNA mismatches on phage immunity provided by Cas12a, we developed a heterologous type V-A CRISPR-Cas12a system in *Escherichia coli*. We expressed Cas12a from *Francisella novicida* and various pre-crRNAs in *E. coli* K12 and infected these cells with lambda phage to measure the immunity provided by Cas12a (Fig. 1a). We used λ_vir_, a mutant of lambda phage that cannot engage in lysogeny and is locked into the lytic mode of replication. This eliminates CRISPR self targeting that could occur if a target phage becomes a lysogen in the bacterial genome. Cas12a was expressed either from a relatively weak constitutive promoter^49, 63^ or a relatively strong inducible promoter (P_BAD_). We chose two lambda genomic targets: one target was in an intergenic region upstream of gene J and the other target was inside the coding region of gene L (Fig. 1a). The Cas12a expression system exhibited a high level of protection for both promoters, with targeting crRNAs showing about 10^6^ fold less phage infection than the non-targeting control (Fig. 1b).

We designed 4 mutant crRNAs with varying levels of in vitro cleavage defects (Fig. 1c) and tested their effects on phage defense (Fig. 1b). These mismatches spanned the target with one in the seed region, one in the mid-target region, and two in the PAM-distal region. We observed a strong defect for the seed mutant when we used the weaker promoter to express Cas12a. However, this defect was reduced upon Cas12a overexpression using the stronger promoter (Fig. 1b), consistent with the defect being caused by reduced Cas12a targeting. Mid-target and PAM distal mismatches caused almost no visible defects in protection for the gene J target and small defects for the gene L target when Cas12a expression was controlled by the stronger promoter. These results correlated with the cleavage defects measured in vitro for the corresponding mismatched crRNAs (Fig. 1c, Supplementary Fig. 1).

We then tested the effects of mismatched crRNAs in liquid culture when Cas12a was expressed from the stronger promoter. Cells containing a matching crRNA grew at the same rate as cells that were uninfected with phage, demonstrating complete Cas12a protection in the time frame tested (Fig. 1d). In contrast, most mismatched crRNAs caused lysis to occur regardless of the mismatch location in the target. For the gene J target, a crRNA mismatch in the seed region caused lysis to begin 1 h after infection, similar to a culture bearing a non-targeting crRNA. This indicated that the seed mismatch was allowing nearly full phage escape, consistent with this mismatch causing the largest reduction of target cleavage in vitro (Fig. 1c).

In contrast, the seed mismatched crRNA against gene L provided protection for several hours post infection, with lysis beginning 3 h post-infection (Fig. 1d). Mismatches in the mid- or PAM distal region offered protection until four or five hours following infection. Interestingly, the rate of cleavage for these crRNAs did not always correlate with the level of protection provided in liquid culture (Fig. 1c-d). Some crRNA mismatches that caused small decreases or no significant effect on cleavage rates in vitro led to lysis of the culture (e.g. gene J position 8 and gene L position 15). These results indicate that loss of cleavage caused by crRNA mismatches did not completely account for loss of immunity.

**Figure 1.**
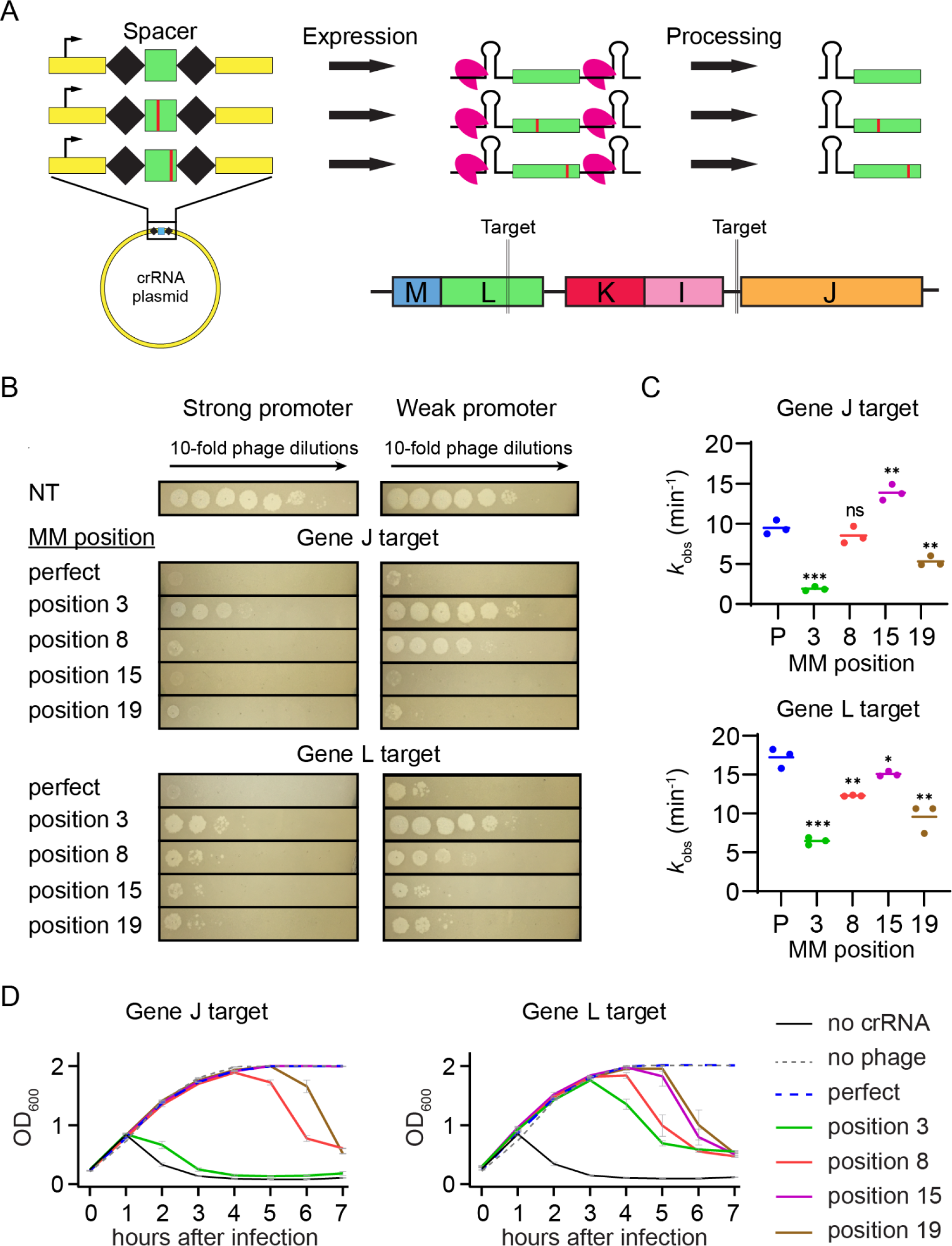
Effects of mismatched crRNAs on Cas12a-mediated phage defense. (A) Schematic of crRNA expression and processing by FnCas12a and crRNA phage target locations. crRNA mismatches were introduced by mutating individual nucleotides in the spacer sequence. After expression of the pre-crRNA, Cas12a processes it into a guiding crRNA that partially matches the lambda phage genome targets upstream of gene J and in the coding region of gene L. See Supplementary Fig. 1a for target and crRNA spacer sequences. (B) Measurement of phage protection provided by crRNAs with and without target mismatches. Spot assays were performed with bacteria expressing FnCas12a and a crRNA construct that either perfectly matches the lambda phage genome (perfect) or has a crRNA mismatch (MM) at a position in the spacer (position x, sequences shown in Supplemental Fig. 1a). A non-targeting crRNA construct (NT) was used as a negative control. Lambda phage was spotted on cells with 10-fold decreasing concentration at each spot going from left to right. Expression of FnCas12a and pre-crRNAs were controlled by a stronger inducible P_BAD_ promoter or a weaker constitutive promoter. (C) Observed rate constants for in vitro cleavage by Cas12a armed with crRNAs containing target mismatches. Plasmids bearing target sequences for gene J or L were used to measure Cas12a cleavage. Mismatch positions or perfect crRNAs (P) are indicated on the horizontal axis. Data from three replicates are plotted. * *P* ≤ 0.05, ** *P* ≤ 0.01, *** *P* ≤ 0.001, ns = no significant difference compared to the perfect crRNA based on unpaired two-tailed t-test. See Supplementary Fig. 1b-c for gels and quantification. (D) Growth curves for *E. coli* expressing mismatched crRNAs following phage infection. Bacteria containing the P_BAD_ FnCas12a expression plasmid and various crRNA expression plasmids were inoculated in liquid culture and induced immediately. Lambda phage was added 1.5 hours after inoculation and OD_600_ measurements were taken every hour. Bacteria expressed no cRNA, a crRNA with no mismatches to the target (perfect) or a crRNA with a mismatch at the indicated position (position x). A no phage condition was performed as a negative control. The average of two replicates is plotted, with error bars representing standard deviation.

### Phage target mutations depend on location of existing mismatches

Our initial results showed that crRNA mismatches have less of an effect on solid media than in liquid culture. We hypothesized that these differences were caused by phage mutation upon CRISPR immune pressure. Unlike on solid medium, phage mutants that arise can quickly and uniformly spread throughout the culture in a liquid medium. Thus, pre-existing mismatches or mismatches that arise through imperfect DNA repair following Cas12a cleavage may accelerate the selection for escape mutants as they quickly spread throughout the population, causing lysis in liquid culture. These results overall suggested that loss of protection from crRNA mismatches is due in part to emergence of phage mutants that further disable CRISPR interference.

To test this hypothesis, we investigated mutations that arose in phage populations in response to CRISPR pressure by Cas12a effector complexes with or without pre-existing crRNA mismatches (Fig. 2a). We isolated phage from liquid cultures of *E. coli* expressing matching or mismatched crRNAs and PCR amplified the regions of the phage genome that were being targeted. High-throughput sequencing was then used to identify mutations in the target regions.

We first quantified the percent of the phage population that had mutations in genomic regions targeted by Cas12a over time in liquid culture. Phages targeted by a matching crRNA gradually developed target mutations over time, with about 20% of the phage population becoming mutated after the 8 hour time course (Fig. 2b). A crRNA mismatch at any of the positions we tested led to a large acceleration of mutant emergence causing the phage population to become almost entirely mutated after 4 hours. Interestingly, phages exposed to bacteria expressing crRNAs with a seed mismatch also rapidly mutated, even though our in vitro results showed the original crRNA mismatches were highly deleterious for target cleavage (Fig. 1c). For most individual replicates of our samples, we did not observe substantial variability in the distribution of mutations after the phage population became highly mutated (Supplementary Fig. 2a). We therefore chose to focus on the longest time point (8 h) for further analysis.

As previously shown in other CRISPR systems^10, 35, 64^, phage populations targeted with a perfectly matching crRNA developed mutations in the seed region of the target (Fig. 2c). When the sequences of the crRNA plasmids were changed to create mismatches between the crRNA and the phage genome target, the position of phage target mutations became substantially more variable. A crRNA mismatch at position 3 for the crRNA targeting the region upstream of gene J caused 9 different mutations to appear at 8 positions spread across the PAM and seed, none at position 3 as expected given the pre-existing mismatch. For the gene L target, a crRNA mismatch at position 3 only caused 2 different mutations to appear, with one of them being the predominant mutation seen when targeting with a matching crRNA. Mismatches in the mid-target region at position 8 also caused seed mutations to arise. Surprisingly, PAM-distal crRNA mismatches caused enrichment of PAM-distal mutations and prevented nearly all seed mutations from emerging. Most of the mutations present in liquid culture were also observed when sequencing phage from spot assays, although the distribution differed in some cases (Supplementary Fig. 2b). This indicates that the differences we observed between our solid media and liquid cultures experiments were caused by the increased mobility of phages in liquid culture, and were unrelated to the types and location of mutations that could arise.

The genomic context of target sequences had a clear effect on the types of mutants that emerged (Fig. 2d, Supplemental Fig. 2c). For the matching crRNA targeting the region upstream of gene J, the most common mutation observed was a single nucleotide deletion at position 6. The most common mutation for the gene L target was a single nucleotide substitution at position 2 which is a wobble base position in the codon. Similar to the other mismatched crRNA constructs targeting gene L, most mutations we observed were either silent or caused amino acid changes from valine, threonine or serine to alanine or from proline to leucine. No deletions were observed in the gene L target in any samples with crRNA mismatches, while deletions were observed in the gene J upstream target in samples with crRNA mismatches at positions 15 and 19. This difference in mutational variability reflects the more vulnerable target region of gene L where base substitutions are likely to change the amino acid sequence of the protein and single deletions will cause frame-shifts. The relatively weak constraints on viable mutations in the upstream region of gene J may enable more routes for escape from Cas12a targeting, resulting in the loss of protection at earlier time points (Fig. 1d).

**Figure 2.**
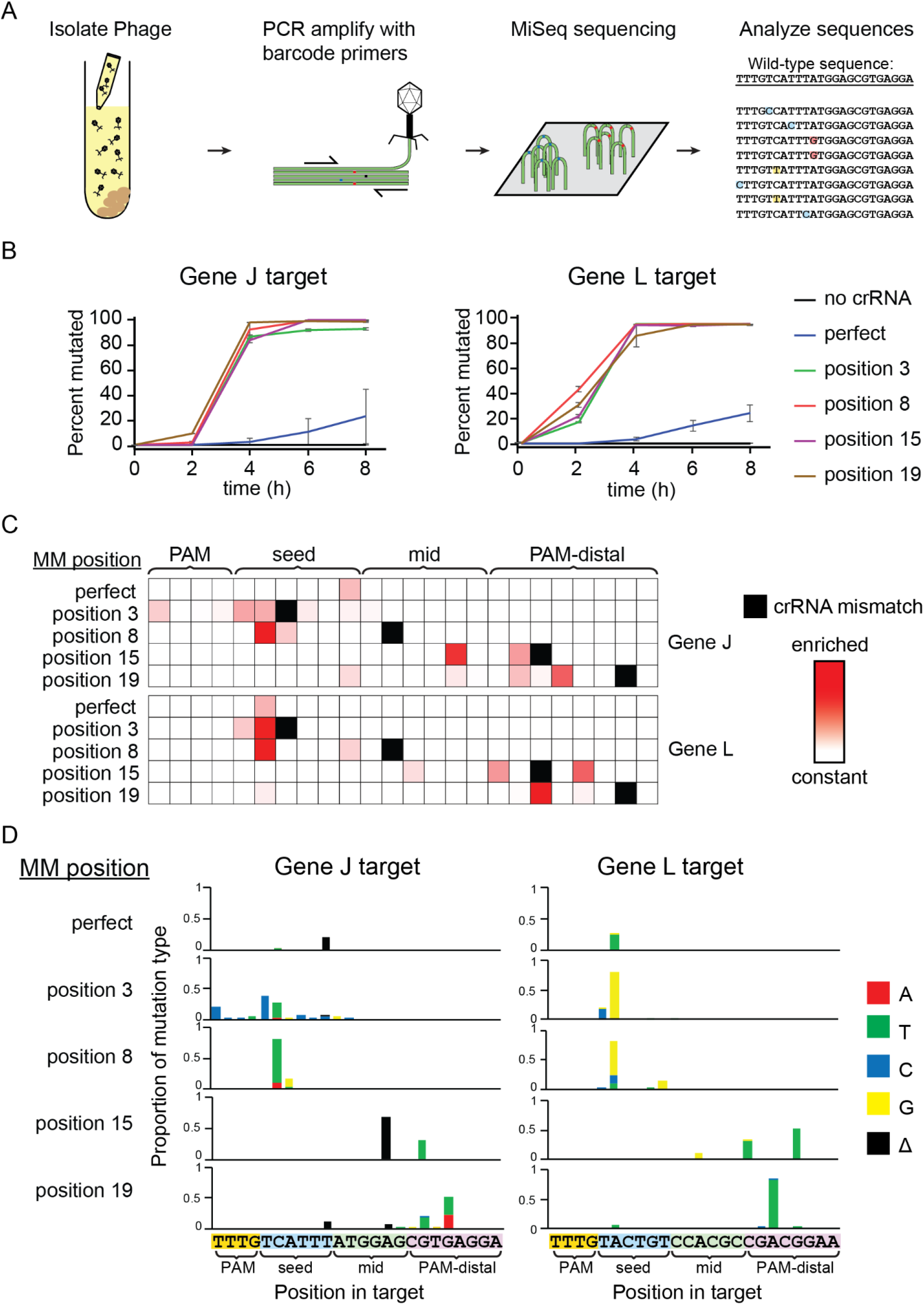
crRNA mismatches cause emergence of diverse lambda phage mutations. (A) Schematic of workflow for determining the genetic diversity of phage exposed to interference by Cas12a. Phage samples were collected from liquid cultures at various time points and the target region was PCR amplified. Mutations were observed using MiSeq high-throughput sequencing of these amplicons. (B) Line graph tracking the fraction of phage with mutated target sequences over time. Samples were taken from liquid cultures at time points after phage infection. The “0 h” samples were taken directly after addition of phage to the culture tubes. The average of two replicates are plotted with error bars representing standard deviation. (C) Heat map showing the location of enriched phage mutations in target regions at the 8 h time point. Z-scores for abundance of single-nucleotide variants, including nucleotide identity changes or deletions, were determined for each sample relative to the non-targeted control phage population. Enriched sequences indicate high Z-scores. Z-scores range from 0 (white) to 7.1 (darkest red). Single nucleotide deletions are shown at adjacent position to the 3′ side. Positions with crRNA mismatches are labeled with solid black boxes in the heat map. (D) Graphs showing single-nucleotide variations for mutated phage target sequences present at the 8 h time point. Bar graph height shows the proportion of sequences in each sample with the mutation type at each position in the target. Deletions (Δ) are plotted at adjacent position to the 3′ side. Values plotted are the average of two replicates. Individual replicates are shown in Supplementary Fig. 2c.

### Multiple mismatches in the PAM distal region allow phage escape from Cas12a

A striking result from our sequencing of mutant phage populations was the emergence of PAM-distal mutants upon challenge with crRNAs containing PAM-distal mismatches. Given that seed mutants appeared when other Cas12a crRNAs were used, these results suggested that multiple PAM-distal mismatches are at least as deleterious for Cas12a cleavage as a seed mismatch combined with a PAM-distal mismatch. It has been shown that pairs of mismatches are more deleterious for Cas12a cleavage when they are closer together, including when both are in the PAM-distal region^53, 56^. Together, these results suggest that double mismatches in the PAM-distal region can lead to phage escape from Cas12a.

To test this, we added second PAM-distal crRNA mismatches to crRNAs targeting gene J that initially contained a single PAM-distal mismatch. We chose the second mismatch position based on phage mutants that appeared when exposed to the original mismatched crRNA (Fig. 2d, Supplementary Fig. 3a). Adding a second mismatch at position 14 to the crRNA that contained a mismatch at position 15 caused a small defect in phage protection (Fig. 3a). Although this mismatch pair had small effects on the overall cleavage rate in vitro regardless of the mismatch type at position 14 (Fig. 3b, Supplemental Fig. 3b-c), we did observe a cleavage defect, in which the DNA was nicked by Cas12a through cleavage of only one strand (Supplemental Fig. 3b). This defect in second-strand cleavage may allow more phage infection, resulting in partial loss of phage defense on solid media (Fig. 3a). Consistently, bacteria expressing a crRNA with a position 15 mismatch did not lyse in liquid culture (Fig. 1d), despite the emergence of the position 14 mutation (Fig. 2c-d). In contrast, adding any type of second mismatch at position 16 to a crRNA that contained the original mismatch at position 19 caused nearly complete loss of protection (Fig. 3a) consistent with lysis observed for the position 19 mismatched crRNA in liquid culture (Fig. 1d) and the dramatic loss of cleavage activity in vitro (Fig. 3b, Supplementary Fig. 3b-c).

**Figure 3.**
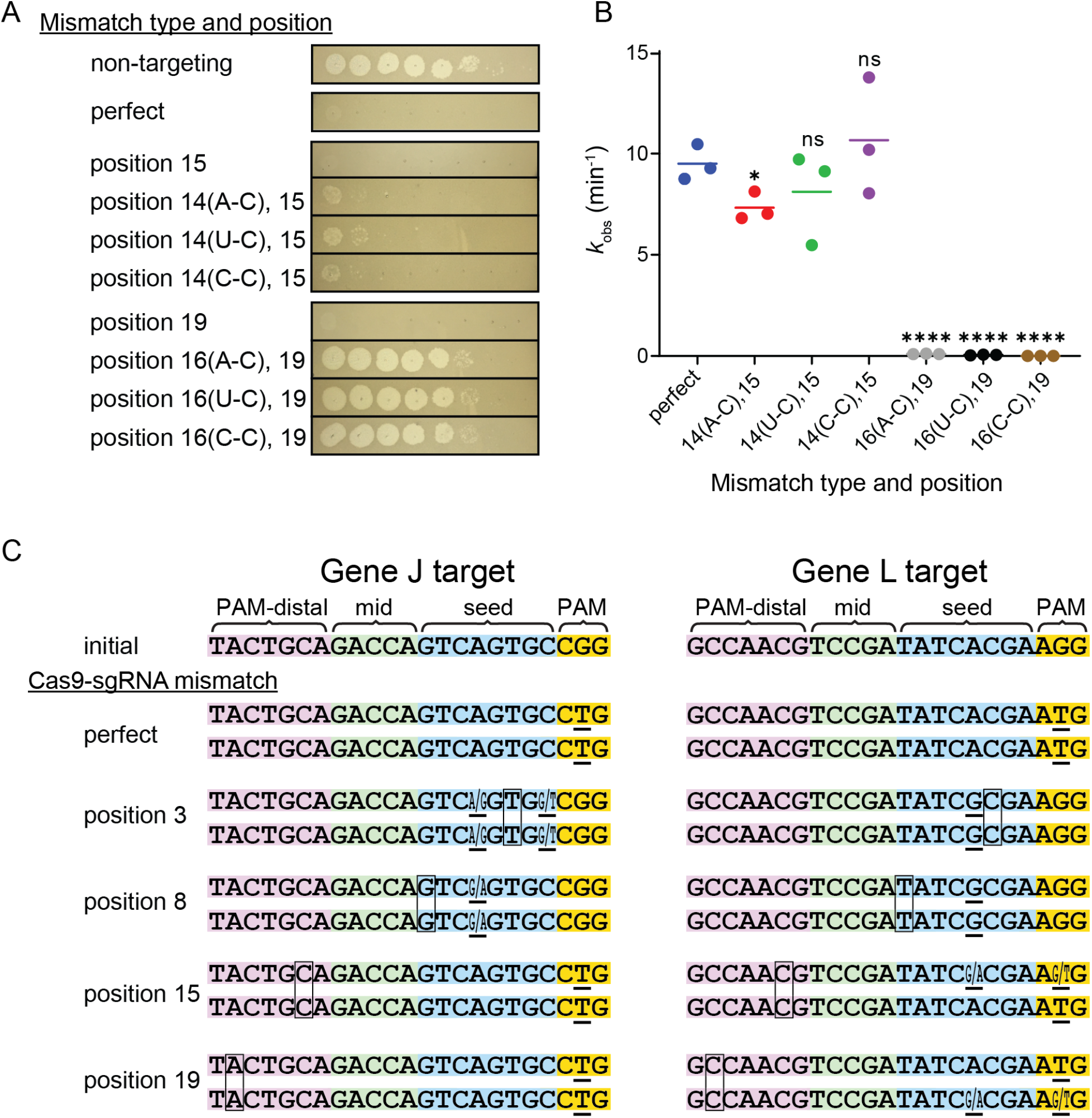
Effects of PAM distal mismatches on Cas12a and Cas9 cleavage and phage protection. (A) Spot assays performed using *E. coli* expressing FnCas12a and a crRNA that perfectly matches the lambda phage gene J target (perfect) or has mismatches at the indicated positions. Three types of second mismatches were added and the type of the mismatch is indicated in parenthesis next to the position number. See Supplementary Fig. 3a for crRNA spacer sequences. (B) Observed rate constants for in vitro cleavage by Cas12a armed with crRNAs containing two target mismatches. Cleavage was measured for plasmid DNA containing a gene J target. The types of mismatches for the second mismatch are indicated. * *P* ≤ 0.05, **** *P* ≤ 0.0001, ns = no significant difference compared to the perfectly matching crRNA based on unpaired two-tailed t-test. See Supplementary Fig. 3b-c for gels and fit data. (C) Sequences of target regions in the lambda phage genome after exposure to SpCas9 armed with mismatched sgRNAs. sgRNAs contained mismatches at the indicated position or matched the target (perfect). Mismatch locations in sgRNAs are indicated with a box. Mutations relative to the initial target sequence are indicated with an underline. Positions with ambiguous base calls are indicated with two bases (X/Y) at that position. Target sequences were interpreted from Sanger sequencing chromatograms (see Supplementary Fig. 4c).

Our results show that some pairs of PAM-distal mismatches are deleterious enough to cause escape from CRISPR-Cas12a immunity. We wondered whether other Cas effectors are subject to a similar route of escape. To determine whether PAM distal mutants could also arise against another Cas effector, we performed the analogous experiments using *Streptococcus pyogenes* Cas9 (SpCas9) with single guide RNA (sgRNA) mismatches at the same positions tested for FnCas12a.

Similar to FnCas12a, sgRNAs containing mismatches caused minimal defects in SpCas9-mediated phage defense on solid media but accelerated lysis in liquid culture (Supplementary Fig. 4a-b). To determine whether lysis occurred due to the emergence of phage mutants, we PCR amplified the target regions of phage collected from these lysates and sequenced the amplicons using Sanger sequencing. Although mixed populations were observed using this method, phage target mutations were clearly visible only in the PAM and seed region of Sanger sequencing chromatograms for all sgRNAs tested with and without mismatches (Fig. 3c, Supplementary Fig. 4c). These results suggest that the sensitivity to multiple PAM distal mismatches is not common to all CRISPR effectors, and that Cas12a may be more susceptible to phage escape via PAM distal mismatches than Cas9.

### Phage targeted with mismatched spacers develop conditional escape mutations

Our results suggest that individual mismatches are often not sufficiently deleterious to allow phages to escape Cas12a targeting. Instead, the combination of the pre-existing mismatch and an additional mutation in the target is necessary for escape to occur. It remains unclear to what extent these new mutations contribute to phage escape in the presence of a pre-existing mismatch. We chose to pursue further experiments using the crRNA with a seed mismatch targeting gene J because although it was highly deleterious for cleavage in vitro (Fig. 1c), it caused rapid phage mutation in liquid culture (Fig. 2b). In addition, this mismatch caused the largest variety of mutants to arise for all the crRNAs we tested with mutations at nearly all positions in the seed region (Fig. 2d). We hypothesized that this target in an intergenic region was less restrictive of mutation, exacerbating the defect of this crRNA mismatch in vivo.

To test this hypothesis, we generated mutated phage populations using the seed mismatched crRNA targeting gene J. We first infected *E. coli* cells expressing the mismatched crRNA with lambda phage in liquid culture at a wide range of MOIs. The phages were able to clear the culture at MOIs greater than 1.5 x 10^-3^ (Supplementary Fig. 5a). We then analyzed the genomic diversity of the phage population in the targeted region using high-throughput sequencing. The phage population retained the wild-type sequence of the target region at the two highest MOIs tested (0.15 and 0.075 MOI), indicating that the wild-type phage can overcome Cas12a-mediated immunity when the bacteria are exposed to enough phage particles (Fig. 4a). At the lowest MOIs tested, 1.5 x 10^-4^ and 1.5 x 10^-5^, 99% of the phage population contained a single mutation at the first position of the protospacer. Mutations that arose were most varied at intermediate MOIs. These mutations were in the seed region or mid target region near the existing crRNA mismatch. The number of different mutations observed was also higher compared to the bacterial strain with a matching crRNA to the WT phage target.

Next, we harvested phage from the cultures at all of the MOIs tested and compared protection against this mutant phage population by a crRNA that perfectly matched the wild-type target and a crRNA bearing the original seed mismatch used to generate the mutant population. Visible infection using these new phage lysates was first observed when using phage that was 48% mutated (Fig. 4b, Supplementary Fig. 5b). While the perfect crRNA still offered some level of protection against the mutated phage, the crRNA containing the mismatch resulted in complete loss of protection (Fig. 4c). These results indicate that some mutations that emerge in the phage population are only significantly deleterious to Cas12a interference in the presence of the pre-existing mismatch, revealing the importance of combined mismatches for phage escape.

**Figure 4.**
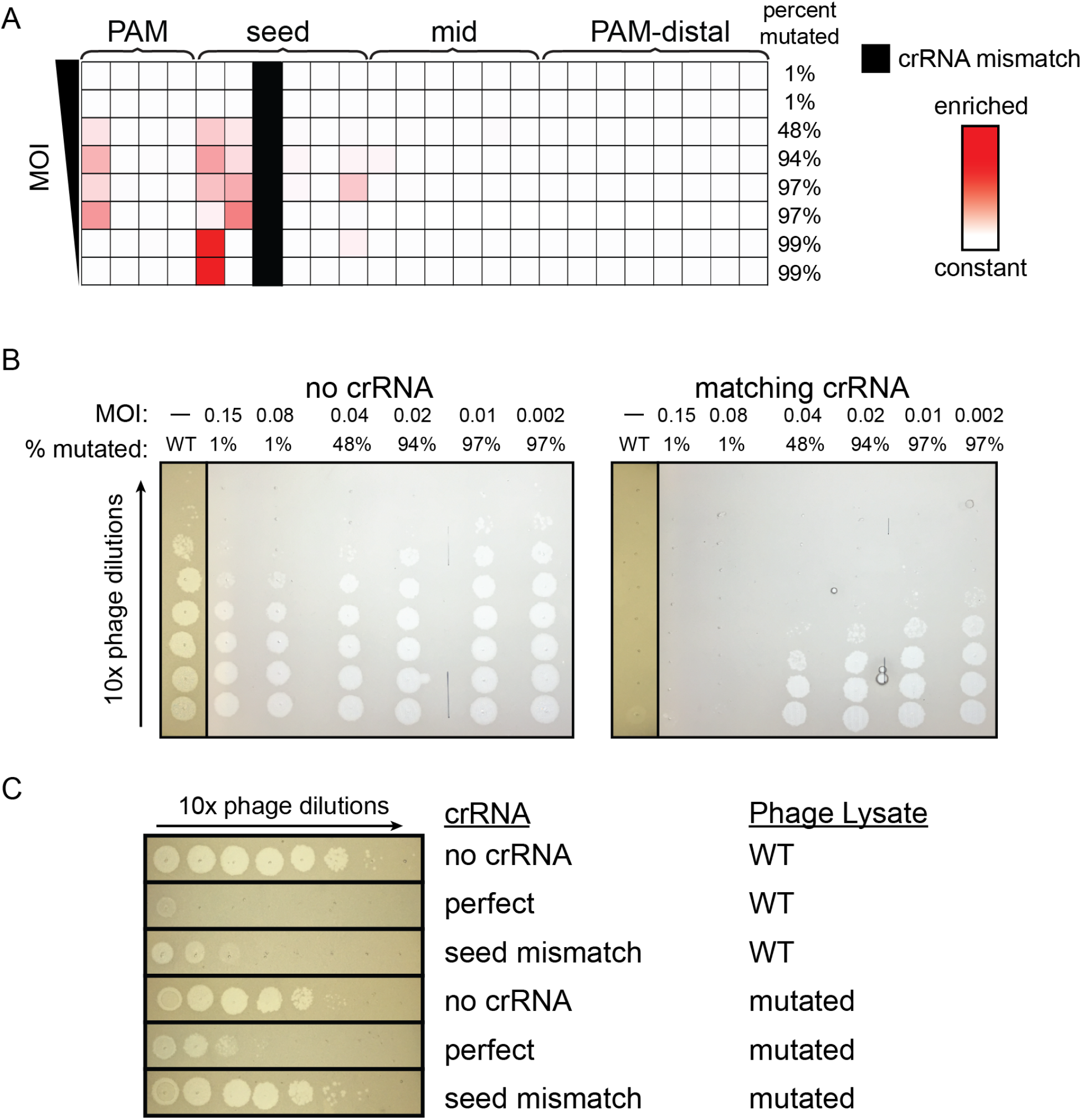
Combined mismatches are necessary for complete phage escape. (A) Heat map showing the position of phage mutations that arose when infecting bacteria expressing seed mismatch crRNA at different MOIs. Phage was harvested from liquid cultures containing bacteria expressing FnCas12a and a crRNA with a C-T mismatch at position 3. Phage was added to the culture at mid-log phase at a range of MOIs starting at 0.15 and serial 2-fold dilutions from 1/2 to 1/32 and an additional sample at an MOI of 1.5 x 10^-3^. Phage was harvested 5 hours after infection. High-throughput sequencing was used to determine the percent of phages in each that had a mutation in the target region. The heat map shows the positions in the target that were enriched with mutations. These positions are colored darker red according to their Z-score relative to the control phage population. Z-scores range from 0 (white) to 10.1 (darkest red). The position of the initial crRNA mismatch is indicated in solid black (B) Spot assays using phage isolated from liquid cultures as described in (A) on bacteria expressing a matching crRNA. Phage harvested in (A) was 10-fold serial diluted and spotted on bacteria with a crRNA matching the wild-type lambda phage genome target (matching crRNA) or bacteria without a crRNA guiding Cas12a (no crRNA). Wild-type phage controls were spotted on these same bacterial strains. Phages harvested from the lowest MOI cultures were omitted due to their low titer which prevented visible plaque formation on the CRISPR active *E. coli* strain. See Supplementary Fig. 5b for full plates. (C) Spot assays using mutationally diverse phage on bacteria expressing crRNAs with and without mismatches. Phage was isolated from the liquid culture as described in (A) that was initially infected with phage diluted 1:8. Mutated phage and unmutated control phage (WT) were then used for spot assays on bacterial lawns expressing FnCas12a and a matching crRNA (perfect), a crRNA with the original seed mismatch, or no crRNA as negative control.

### Combining mismatched spacers increases level of protection

Our results indicated that loss of protection by Cas12a due to crRNA mismatches was only partially caused by loss of Cas12a cleavage due to the pre-existing mismatch, and that mutant emergence generating a second mismatch also contributed substantially to this loss of protection. These results imply that Cas12a mismatch tolerance should enable stronger and longer term protection under conditions where phage mutants are less likely to emerge. To further test this, we introduced both the gene J and L crRNAs into a CRISPR array for co-expression of both crRNAs. It has been demonstrated previously that bacterial populations with diverse spacer content in their CRISPR arrays show robust immunity against phage at the population level as phage are unable to develop escape mutations in all of their targeted genomic regions^65, 66^. Cas12a expressed along with multiple different cRNAs shows increased phage resistance compared to when a single crRNA is present, but the mechanism of this increased resistance is not clear^64^. We investigated this mechanism further in the context of our previous experiments with mismatched crRNAs. If the loss of protection due to a crRNA mismatch is caused only by a slowing of the rate of cleavage, then two different mismatched spacers should not provide more protection than one spacer repeated twice. Alternatively, if phage mutant emergence significantly contributes to loss of protection in the presence of a crRNA mismatch, two different mismatched spacers should provide better protection than a single mismatched spacer repeated twice.

Consistent with the second possibility, the CRISPR construct with two unique mismatched spacers (hereafter referred to as double spacer construct) showed a significantly higher level of protection than either of the crRNA constructs with two copies of a single mismatched spacer (hereafter referred to as single spacer construct) when measured by plaque assay (Fig. 5a). Similar to liquid cultures with bacteria expressing a single copy of the crRNA, we observed faster lysis of the gene J targeting crRNA in comparison to the gene L targeting crRNA, consistent with the higher chance of escape mutant emergence against the gene J crRNA. Bacteria expressing the double spacer construct showed slowed growth between 1 and 2 hours but recovered quickly and did not lyse over the time course tested (Fig. 5b). Phage from these cultures was harvested over time and used to infect CRISPR inactive bacteria to determine the relative titers. Phage titers decreased over time in cultures expressing the double spacer construct, while the phage titer increased over time in cultures with cells expressing the single spacer constructs (Fig. 5c).

Spotting these same phage lysates on CRISPR active cells showed no noticeable infection by lysate harvested from the double spacer culture, but moderate infection by the single spacer lysate (Supplementary Fig. 6a), suggesting that escape mutants did not emerge from bacteria expressing two different mismatched crRNAs. Consistently, sequencing of both target regions in individual plaques revealed mutations in only one of the two target regions (Fig. 5d, Supplementary Fig. 6b). Together with our previous results, these results suggest that loss of protection due to a crRNA mismatch is caused by a combination of loss of Cas12a targeting and the emergence of mutant phages that further block CRISPR interference.

**Figure 5.**
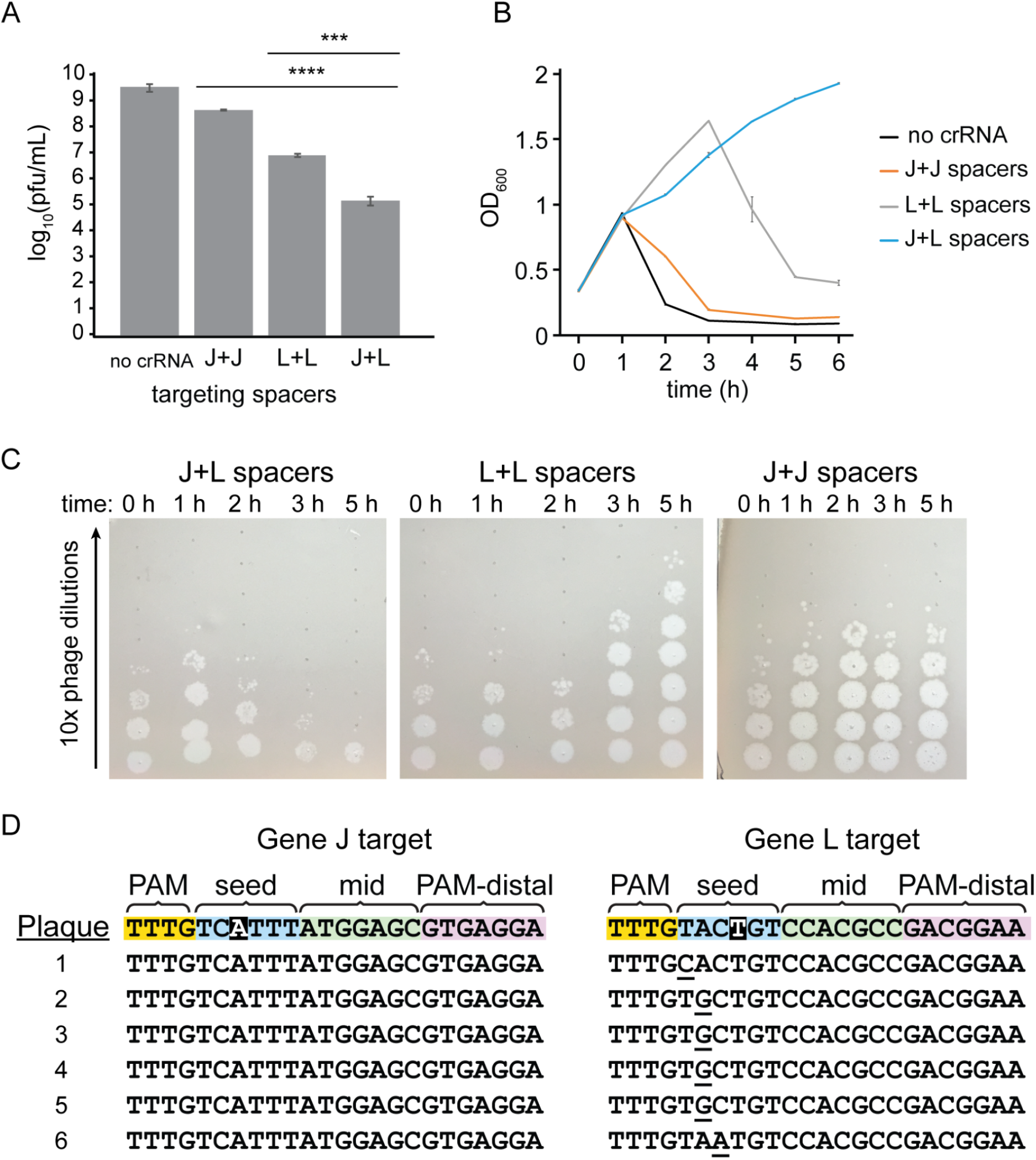
Multiple mismatched crRNAs provide more protection than individual mismatched crRNAs. (A) Number of plaques formed on lawns of bacteria expressing multiple mismatched crRNAs. Plaque assays were performed using bacteria containing a plasmid expressing FnCas12a along with different crRNA expression plasmids. Each crRNA expression plasmid contains two spacers: both targeting gene J (J+J) both targeting gene L (L+L) and one spacer targeting each of gene J and gene L (J+L) or no crRNA. Plaque forming units (pfu) was calculated using the number of plaques on each plate and the volume of phage lysate added. The average of multiple replicates (n = 2 to 4) is plotted with error bars representing standard deviation. Unpaired, two-tailed t-tests were used to determine the statistical significance of each of the single spacer constructs compared to the double spacer construct, *** *P* ≤ 0.001, **** *P* ≤ 0.0001. (B) Growth curves using the same bacterial strains described in (A). Cultures were grown to ∼0.25 OD_600_ in induction media and infected with lambda phage at the “0” hour time point. OD was then measured every hour for 6 hours. (C) Spot assays used to measure the titer of phage over time in phage infection cultures. Spot assays were performed using 10-fold serial dilutions of phage harvested from cultures in (B) that infected bacterial strains with two mismatched spacers at different time points on lawns of CRISPR-inactive *E. coli*. (D) Sequences of both CRISPR targets in single phage plaques for phage harvested from *E. coli* cultures expressing a double spacer construct. The two crRNAs contained mismatches at positions highlighted in black. Target regions for the gene J and gene L target were sequenced for six individual plaques using Sanger sequencing. Target sequences are aligned to the WT sequence (top row) and mutations are underlined. See Supplementary Fig. 6b for chromatograms.

### Pre-existing target mutations cause different CRISPR escape outcomes

We have shown that target mismatches artificially introduced by changing crRNA sequences accelerate phage escape and increase the diversity of mutations that appear. We wanted to determine if the same effect would appear if the crRNA-target mismatch was instead caused by a phage genome mutation. We first generated clonal phage populations with single target mutations by isolating individual plaques of mutant phage that emerged following exposure to Cas12a bearing various crRNAs (Fig. 6a). Using a crRNA containing a seed mismatch, we isolated phages with mutations in the PAM (T-2C) or seed (C2A) (Supplementary Fig. 7a,b), while a crRNA containing a mismatch at position 19 allowed us to isolate two separate plaques containing phage with a mutation at position 16 (G16T) (Supplementary Fig. 7c,d). After propagating phage from these plaques, we challenged the mutant phages to CRISPR pressure by bacteria expressing crRNAs with a spacer matching the wild-type phage genome target. We observed that the phage with a seed region mutation caused rapid lysis of CRISPR active bacteria (Fig. 6b). This indicated that the C2A mutation was a complete escape mutation. However, phage mutations in the PAM or PAM-distal region caused delayed lysis to occur. In particular, of the two G16T isolates, only one caused lysis to occur in some of the experimental replicates (Fig. 6b, data not shown for isolate where lysis did not occur).

We harvested phage from the previous cultures and sequenced PCR amplicons of the phage genome targets using Sanger sequencing. In the seed mutant (C2A) phage cultures, the phage retained the same seed mutation and did not develop additional mutations (Fig. 6c, Supplementary Fig. 7b), further indicating that C2A is a bona fide escape mutation on its own. In phage with pre-existing mutations in the PAM, mutations appeared at the edge of the seed region (Fig. 6c, Supplementary Fig. 7a). In phage with a pre-existing mutation in the PAM-distal region at position 16, mutations appeared at positions 14 or 18 for phage harvested from cultures that lysed. Repeating the same experiment with PAM-distal mutants in the gene L target similarly caused further mutations to occur in the mid-target and PAM-distal regions (Supplementary Fig. 7e-h).

We proceeded with further experiments using only replicates in which a clonal phage population was generated based on an unambiguous Sanger sequencing chromatogram (Supplementary Fig. 7a,b,d). Using these phage, we sought to verify that these second mutations were allowing CRISPR escape. We compared infection of bacteria expressing the matching crRNA by purified phage containing a single target mutation and phage with two target mutations. As expected, phage with the seed target mutation infected bacteria expressing the perfect crRNA at the same level as bacteria expressing a non-targeting crRNA (Fig. 6d). Phage with single target mutations in the PAM or PAM distal region infected bacteria expressing the perfect cRNA at a level close to wild-type phage, while phage with a second mutation infected 10^4^ to 10^5^ times more (Fig. 6d). We conclude that target mutations that do not lead to significant CRISPR escape can accelerate the appearance of second mutations that allow complete escape.

**Figure 6.**
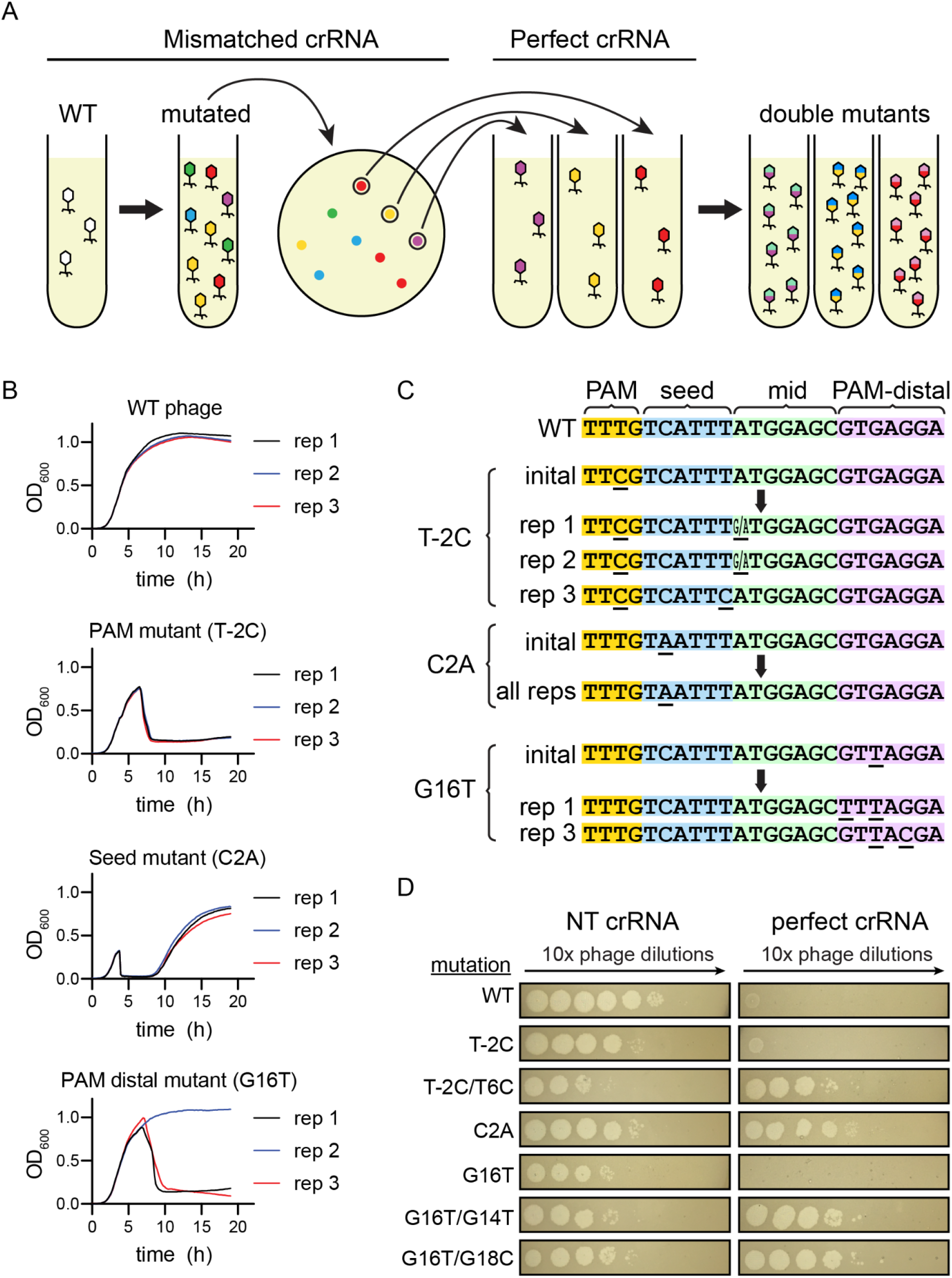
Generation of double mutant phage is driven by insufficiently deleterious mutations. (A) Schematic of the process for generating and purifying single mutant phage populations. Wild-type phage was used to challenge *E. coli* expressing a crRNA with a mismatch to the target in the phage genome in liquid culture. The resulting phage were isolated and used for a plaque assay on lawns of bacteria expressing the same mismatched crRNA. Single plaques were isolated and again used to challenge bacteria expressing a mismatched crRNA in liquid culture, further purifying and propagating single mutants. Finally, single mutant phages were used to challenge bacteria expressing a perfectly matching crRNA in liquid culture to determine whether second mutations would appear. (B) Growth curves of bacteria expressing a perfectly matching crRNA challenged with wild type phage and phage with various single target mutations. Locations of the single mutations in the target are labeled (PAM mutant, Seed mutant, and PAM distal mutant). Position and type of mutations are indicated in parenthesis. Three biological replicates are shown separately for each experimental condition. (C) Diagram of initial and selected mutations that appeared when a single mutant phage was used to challenge bacteria expressing a perfectly matching crRNA. Initial mutants are single mutants that were generated and purified as described in (A). Sequences below arrows show phage mutants that appeared in different biological replicates (rep 1, 2 or 3) after initial mutant phage lysates were used to infect bacteria expressing a perfectly matching crRNA in liquid culture. Positions with ambiguous base calls are indicated with two bases (X/Y) at that position. Target sequences were interpreted from Sanger sequencing chromatograms (see Supplementary Fig. 7). (D) Spot assays challenging bacteria expressing a perfectly matching crRNA with various single and double mutant phage lysates. WT phage or phages with the indicated target mutations were spotted on bacteria expressing a non-targeting crRNA (left column) and a perfectly matching crRNA (right column).

### Large deletions pre-exist in non-essential regions of the lambda phage genome

Our results indicate that mutations can arise rapidly in regions targeted by Cas12a when a pre-existing mismatch is present between the crRNA and target. Previous studies have suggested that DNA cleavage by Cas9 or Cas12a may induce mutation in the target site through DNA repair^35, 38^. Alternatively, it is possible that Cas12a targeting selects mutant phages that are present in the population at the time of infection. To distinguish these two possibilities, we determined if the mutated phage we observed in our CRISPR active samples were present in negative control samples. Control samples were grown under the same conditions as experimental samples but contained a non-targeting crRNA. Thus, phages isolated from these cultures should represent the wild-type lambda phage population, including pre-existing mutants present in the population.

We first examined all of the single nucleotide polymorphisms (SNPs) that became enriched (greater than 1% of reads) in CRISPR active samples. All of the enriched SNPs were found in our control samples, but at low enough levels that we could not rule out that the reads were due to sequencing or PCR error (Supplementary Fig. 8). We then examined single deletion sequences. Two single deletion sequences in the gene J target became enriched and these were consistently present in control samples (Supplementary Fig. 9), indicating that they were pre-existing in the phage population.

Our high-throughput sequencing data suggests that some CRISPR escape mutants are present in the natural phage population, but we also found evidence of mutant generation upon CRISPR challenge. Challenging single mutant phage with a perfectly matching crRNA led to the appearance of second target mutations that allowed escape from Cas12a interference. Although a variety of double mutant phage were generated from our clonal single mutant phage populations, the initial mutation always remained when other mutations appeared (Fig. 6, Supplementary Fig. 7). This indicates that the double mutant phage were not pre-existing in the population and the second mutations may have been generated through recombination or repair after target cleavage.

Our mixed results led us to find other ways of detecting diversity in phage genomes in a population. Large deletions have been previously observed in non-essential regions of the genome of bacteriophage following CRISPR interference^38^. We next used long-read sequencing to determine whether such deletions are pre-existing in our lambda phage population or arise following Cas12a cleavage.

We first confirmed that Cas12a targeting in non-essential, but not essential, genes resulted in large genomic deletions. PCR amplification of the surrounding area of non-essential targets targeted by Cas12a showed a ladder of smaller PCR products (Fig. 7a). Gel purification and sequencing of the smaller PCR products showed that deletions were enriched in the lambda phage genome that removed the protospacer completely (Supplementary Fig. 10). Short homology was observed at each end of the deletion sites, as has been previously observed^38^.

To determine whether the deletions observed following Cas12a challenge were present in the initial population, we used PacBio long-read sequencing to sequence PCR amplicons of the non-essential regions in our lambda phage population unexposed to CRISPR targeting. We found small amounts of deletions in both target regions, which represented less than 0.1% of total sequences. Deletions were both longer and more common in the nin204 region than in the nin146 region. Importantly, deletions that were enriched due to CRISPR interference were also the most abundant deletions found in the pre-existing population (Fig. 7b). These results show that mutant phages existed in the natural population and suggest that these mutants became enriched in our experiments due to a selective advantage in the face of interference by Cas12a.

**Figure 7.**
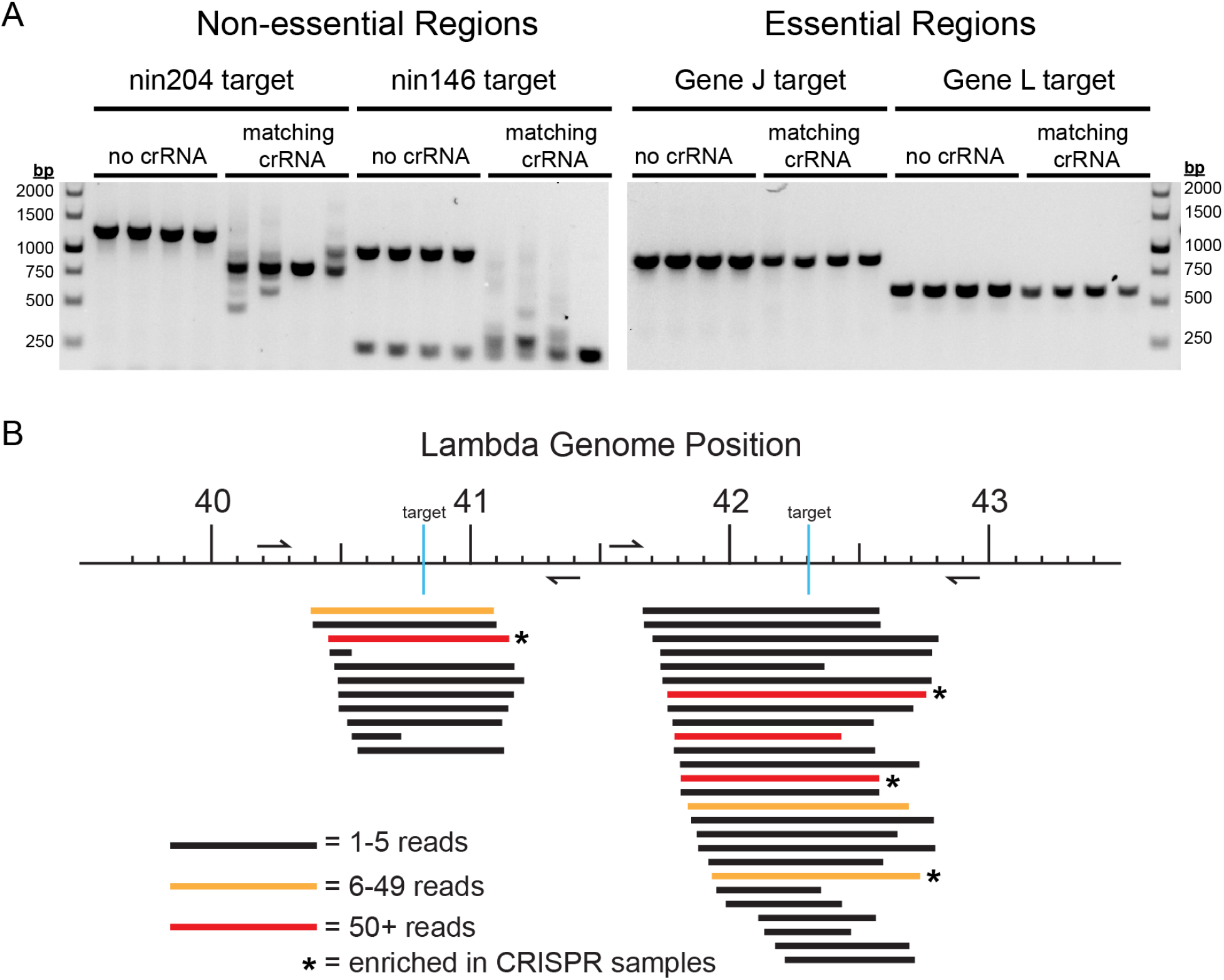
Phage with pre-existing deletions in non-essential genomic regions are selected following Cas12a targeting. (A) PCR amplification of regions surrounding essential and non-essential genes targeted by Cas12a. Spot assays were performed using *E. coli* expressing crRNAs that match two non-essential (nin204 and nin146) and two essential regions (gene J and gene L) of the lambda phage genome. Controls were performed with a plasmid not encoding a crRNA. Phages were isolated from the phage spots and target and flanking regions of the phage genome were PCR amplified and run on an agarose gel. (B) Map of genomic deletions observed by PacBio sequencing of PCR amplicons from phage unexposed to CRISPR targeting. DNA from lambda phage unexposed to CRISPR targeting was used as a template for PCR reactions that amplified the same non-essential regions as in (a). This PCR product was sequenced with PacBio long-read sequencing and the obtained sequences were matched with the wild-type lambda genome sequence to identify any deletions present. These deletions are plotted on the chart relative to their position in the genome. The quantity of each deletion is identified by a color code. Deletions found by Sanger sequencing in CRISPR active samples are indicated with an asterisk (*). See supplementary Fig. 10 for Sanger sequencing chromatograms of PCR amplicons.

## Discussion

In order for Cas12a to be an effective immune effector, it must provide immunity from bacteriophage in diverse conditions. The physical environment controls which bacteria are exposed to which phages, and the dispersal of new phage particles after infection and lysis^9, 67^. We found that Cas12a overall provided more robust immunity on solid media than in liquid culture. The effect of seed crRNA mismatches using either media correlated with the deleterious effect of the crRNA mismatch on rate of cleavage in vitro. Mid-target and PAM-distal mismatches, however, showed a much more drastic effect in liquid culture than defects observed in vitro or on solid media, causing eventual lysis of the bacterial population, sometimes at a rate similar to seed mismatches. These results strongly indicate that the effect of crRNA mismatches varies depending on the environment where phage exposure may occur.

Although phage can escape CRISPR interference with a single mutation in the seed region^10, 35^, our results indicate that a seed mutant is sometimes not deleterious enough to allow complete escape. Instead, the mechanism of phage escape occurs through the emergence of mutations that further weaken CRISPR interference when a mismatch is present. This is supported by the rapid emergence of mutant phage we observed even when a highly deleterious seed crRNA mismatch was present. In addition, the number of different mutations that appeared increased when a crRNA mismatch was present, and the distribution of these mutations greatly varied depending on the location of the mismatch. These results suggest that mismatches between the crRNA and target decrease phage protection by broadening the range of mutations that allow escape. This mechanism also explains the deleterious effect of mismatches at some positions outside of the seed region on immunity in liquid culture that does not appear during in vitro cleavage.

In natural settings, spacer-target mismatches are caused by phage genome mutation rather than CRISPR spacer mutation that would change the sequence of the crRNA^12, 68^. Phages with a single target mutation might not be able to completely escape from certain CRISPR spacers, requiring selection of a second target mutation for full evasion of CRISPR interference^36^. This scenario may become more likely if the seed region, where mutations would normally arise, is located in a critical part of the genome where mutations are highly deleterious. Supporting this, we isolated phage with single mutations that did not cause significant CRISPR escape that then developed second mutations that allowed full escape when exposed to the matching crRNA. In addition, CRISPR spacers with multiple mismatches become more abundant over time in natural populations^46^. Thus, the presence of mutations may drive further mutation in CRISPR targets over time.

When a PAM-distal crRNA mismatch or a PAM-distal target mutation was present, mutations arose in close proximity to the pre-existing mismatch. While this result is surprising, given that PAM-distal mismatches are not highly deleterious to Cas12a cleavage, previous studies have indicated that pairs of mismatches are most deleterious when they are in close proximity^53, 56^. PAM-distal crRNA mismatches did not influence the location of phage target mutations under pressure from Cas9. Accordingly, previous work found little to no synergistic effect of pairs of mismatches in the PAM distal region on Cas9 cleavage^69^. The enrichment of PAM-distal mutations after exposure to Cas12a and not Cas9 may also be influenced by their cut sites, which are in the PAM-distal region and downstream of the target for Cas12a and in the seed region for Cas9.

There is evidence that phage escape mutation is accelerated by error prone repair after cleavage by Cas effectors and mediated by phage recombinases, especially when a crRNA provides a low level of protection^38, 39^. Lambda phage recombination enzymes in particular have been shown to increase the appearance of mutated phage in populations under CRISPR pressure^39^. Our results strongly suggest that mutants that arose upon Cas12a challenge pre-existed in the population, especially for mutants involving single nucleotide or long deletions. We speculate that pre-existing mutations may be propagated in the phage population by these same lambda encoded enzymes. Phage with target mutations in their genome that initially survive interference could be used as recombination substrates to pass along that mutation to other phages in the cell. This could account for the rapid increase in mutant phages we observe and the deleterious effect of phage recombination enzyme knockouts^39^.

Overall, our results reveal that mismatches throughout the crRNA-target duplex can drastically decrease protection provided by Cas12a. However, there are fundamental differences between our heterologous system and natural CRISPR-Cas systems. Adaptation is an important part of CRISPR immunity. As phage mutate and become immune to targeting by existing spacers, new spacers from the phage genome can be added to the CRISPR array^11, 70^. It remains to be investigated how mismatched spacers contribute to acquisition of new spacers in type V systems, especially using a primed mechanism as occurs in type I and type II systems.

## Materials and Methods

### Expression plasmid construction

Cas12a and Cas9 expression plasmids (Table S1) were constructed using pACYCDuet-1. A gene expressing FnCas12a or SpCas9 was inserted downstream of a pBAD promoter in pACYCDuet-1 using Gibson assembly. Promoters were replaced with a previously described low copy P4 promoter^49^ using one piece Gibson assembly. crRNA expression plasmids (Table S1) were constructed using pUC19. A pBAD promoter was inserted into pUC19 in the multiple cloning site with Gibson assembly. “Round the horn” PCR and ligation was used to add a mini CRISPR array with one or two spacers downstream of the pBAD promoter. The same method was used to replace mini CRISPR arrays with Cas9 sgRNA expression constructs.

### Bacterial and phage strains

*E. coli* strain BW25113, a K12 derivative^71^ was used for all experiments along with a modified lambda phage strain (λ_vir_)^72^ that is locked into the lytic lifestyle.

For all CRISPR interference assays, bacteria were transformed with Cas12a and crRNA expression plasmids by heat shock. Transformants were plated on LB plates containing ampicillin at 100 µg/mL and chloramphenicol at 25 µg/mL to select for plasmids pUC19 and pACYCDuet-1, respectively.

### Phage spot assays

Overnight cultures were started using *E. coli* transformed with Cas12a and crRNA expression plasmids in LB media with ampicillin and chloramphenicol for selection. The next day, 20 μL of these overnight cultures were used to inoculate 2 mL cultures in LB media containing 100 µg/mL ampicillin, 25 µg/mL chloramphenicol, 20 mM arabinose, and 10 mM MgSO_4_. The cultures were grown in a shaking incubator at 200 rpm 37 °C until OD_600_ 0.4 was reached. 300 µL of cell culture was added to 3 mL 0.7% soft agar containing the same concentrations of ampicillin, chloramphenicol, arabinose and MgSO_4_. The cell-soft agar mixture was vortexed for 5 seconds and spread onto an LB plate containing ampicillin and chloramphenicol. The plate was dried for 5 minutes. Lambda phage suspended in LB media with 10 mM MgSO_4_ was serially diluted in seven steps with a 1:10 dilution each step. 2 μL of each phage dilution was then spotted on top of the soft agar layer and the plate was dried for 10 minutes. Plates were incubated overnight at 30°C.

### Liquid culture phage assays and growth curves

Overnight cultures were started using a single colony of *E. coli* with Cas12a and crRNA expression plasmids in LB media with ampicillin and chloramphenicol added for selection. The next day, these overnight cultures were used to inoculate cultures 1 to 100 in LB media containing 100 µg/mL ampicillin, 25 µg/mL chloramphenicol, 20 mM arabinose, and 10 mM MgSO_4_. The cultures were grown in a shaking incubator at 200 rpm and 37 °C for 1.5 hours until OD_600_ ∼0.25 was reached.

Lambda phage was added at MOI 0.02, or as indicated in figure legends. For growth curves, OD readings were immediately taken after addition of the phage and this value represented the “0 hour” time point. For growth curves shown in Fig. 1d, Fig. 5a, and Supplemental Fig. 5a, OD was measured at 600 nm wavelength every 1 hour in a WPA biowave CD8000 Cell Density Meter if growing in culture tubes. For growth curves shown in Fig. 6b and Supplemental Fig. 4b, 150 μL cultures were grown in a TECAN infinite M Nano+ 96 well plate reader at 280 rpm and 37 °C and OD measurements at 600 nm wavelength were measured every 10 minutes.

### Phage plaque assays

*E. coli* bacterial cultures were prepared and induced the same way as in the phage spot assays. 50 µL of phage diluted between 1:1 and 1:10^6^ in LB media was added to 300 µL of induced cell culture at OD_600_ 0.4. This mixture was then added to 3 mL 0.7% soft agar containing ampicillin, chloramphenicol, arabinose and MgSO_4_ as in the phage spot assays and the mixture was vortexed for 5 seconds and spread onto an LB plate containing ampicillin and chloramphenicol. The plate was dried for 10 minutes and left to incubate at 37 °C overnight. Plaques were counted the next morning.

### In vitro cleavage assays

Cleavage assays were prepared in reaction buffer (20 mM HEPES, pH 7.5, 100 mM KCl, 1 mM MgCl_2_, 1 mM DTT and 5% glycerol) with a final concentration of 50 nM FnCas12a. FnCas12a RNP complex was formed by incubating FnCas12a and crRNA at a 1:1.5 ratio at 37 °C for 10 min. To initiate cleavage, target plasmid was added to a final concentration of 15 ng/µl and a final reaction volume of 100 µl. The reaction was incubated at 37 °C. Aliquots (10 µl) were quenched at 7, 15, 30, 60, 300, 900 and 1800 s by adding 10 µl phenol-chloroform-isoamyl alcohol (25:24:1 v/v, Invitrogen). After phenol-chloroform extraction, DNA products were separated by electrophoresis on a 1% agarose gel, and visualized with SYBR Safe (Invitrogen) staining. The densitometry of individual DNA bands was measured in ImageJ (https://imagej.nih.gov/ij/). The fraction cleaved was determined by dividing the total cleaved DNA (nicked and linearized DNA) by total DNA (nicked, linearized and supercoiled DNA). Fraction cleaved was plotted versus time and fit to a first-order rate equation to determine an observed rate constant for cleavage (*k*_obs_). Rates were measured in triplicate.

### High-throughput sequencing sample preparation

Phage samples were isolated from spots in spot assays at the highest phage dilution in which a cleared spot was observed to ensure a diverse population of mutant phages would be sampled. A 1 mL pipette tip was poked through a phage spot and the agar in the tip is suspended in 30 µL deionized H_2_O in a 1.5 mL microcentrifuge tube and incubated in a 48°C water bath for 20 minutes to melt the agar and dissolve the phage particles. The tubes were vortexed briefly and incubated in the water bath for another 10 minutes. The water/melted agar mixture that contains the phage was removed.

For phage samples from liquid culture, 50 µL of each culture was transferred to a 1.5 mL tube and bacteria were pelleted from the liquid culture by centrifuging at 15,000 rpm for 5 minutes. The supernatant containing phage particles was then removed.

5 µL of phage solution was used as the template for a 25 cycle PCR reaction with primers containing Nextera adapters (Table S1). These PCR products were cleaned up using the Promega Wizard PCR purification kit and used as a template for an 8 cycle PCR reaction to add barcodes for sample identification. Q5 DNA polymerase (New England Biolabs) was used for all adapter and barcode PCR reactions. Samples were pooled and gel purified using the Promega Wizard PCR purification kit. Gel purified samples were then submitted for MiSeq high-throughput sequencing. Conditions for MiSeq runs were Nextera DNA MiSEQ 150-Cycle which included two 75 base pair paired end reads. Adapter PCR primers were designed so both of the paired R1 and R2 reads overlapped in the entire protospacer region including the PAM.

Samples were prepared for PacBio sequencing by 35 cycle PCR amplification of phage samples isolated from liquid culture. PCR products were purified using the Promega Wizard PCR purification kit and submitted for Pacbio sequencing.

### High-throughput sequencing data processing

A script written in Python 3.8 was used to process .fastq data files received from a MiSeq run. The script extracts target region sequences and determines if the target region contains a mutation relative to the wild-type target sequence. Base substitutions and deletions were classified along with the location of the substitution or deletion relative to the PAM sequence of the target. Mutations were also classified based on the type of mutation (A to C for example). A Microsoft Excel sheet was then created with the data using the “Xlsxwriter” Python package^73^. Z-score calculations and heat maps for each sample were created using Microsoft Excel. Z-scores for each position in the phage target regions were calculated using the average proportion of reads with mutations at each position in experimental samples 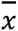 and control samples 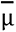 and the standard deviation of the proportion of mutations at all positions in all samples σ.

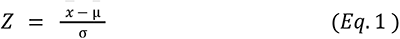

A separate script written in Python 3.8 was used to process data files received from PacBio high-throughput sequencing and find deletions in the lambda phage genome. Sequences were extracted from .fastq files and matched piecewise to the WT sequence of the genome region that was PCR amplified. Deletions are output as coordinates in the PCR amplified region and these coordinates were translated to the lambda phage genome to create the bar graph in Fig. 7b.

Nucleotide diversity was calculated using the proportion of all pairs of sequences *x*_*i*_ and *x*_*j*_ in each sample and the number of nucleotide differences between each pair of sequences π_*ij*_ divided by the length of the PAM and protospacer region (24).

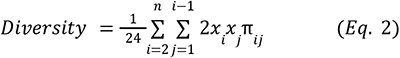

Find the scripts at https://github.com/alexsq2/lambda-phage-CRISPR-mutants

### Sanger sequencing of phage plaques or phage lysate

Phage was isolated from a single plaque by poking into the plaque with a 1 mL pipette tip and the agar was suspended in 20 µL deionized H_2_O in a 1.5 mL microcentrifuge tube and incubated in a 48°C water bath for ∼20 minutes. Melted agar and H_2_O mixture containing phages was transferred to a 1.5 mL microcentrifuge tube.

Phage was also isolated from liquid cultures by transferring 1ml of liquid culture to a 1.5 mL microcentrifuge tube and centrifuging at 15,000 rpm for 5 minutes. Supernatant containing phages was transferred to a clean 1.5 mL microcentrifuge tube.

5 μL of phage solution was used as a template for a PCR reaction that amplifies the target region in the middle of a ∼800 base pair PCR product. One of the primers used for the PCR reaction was used for sequencing of the PCR product.

### Generation and purification of mutant phage

Overnight cultures were started using a single colony of *E. coli* with Cas12a and crRNA expression plasmids in LB media with ampicillin and chloramphenicol added for selection. The next day, these overnight cultures were used to inoculate cultures 1 to 100 in LB media containing 100 µg/mL ampicillin, 25 µg/mL chloramphenicol, 20 mM arabinose, and 10 mM MgSO_4_. The cultures were grown in a shaking incubator at 200 rpm 37 °C for 1.5 hours until OD_600_ ∼0.4 was reached. Lambda phage was added at MOI 0.02. Cultures continued to grow in the shaking incubator for 5 hours. Cultures were transferred to 1.5ml tubes and centrifuged at 5000 rpm for 5 minutes. Supernatant containing phage was transferred to a fresh 1.5 mL tube

Phage was then diluted and used for phage plaque assays on lawns of bacteria expressing the same crRNA as in the previous infection to select against remaining WT phage. Single plaques were isolated and used to infect bacterial cultures again expressing the same crRNA under the same conditions as described above. Cultures were grown in a shaking incubator at 200 rpm 37 °C for 5 hours. 5 µL of phage solution was then used as a template for a 35 cycle PCR reaction with Phusion polymerase to amplify the target region. Sanger sequencing was used to confirm the presence and purity of mutations in the target region.

## Acknowledgements

We thank Michael Baker and Kevin Cavallin of the Iowa State DNA Facility for advice on MiSeq sample preparation and data processing. MiSeq sequencing was performed at the Iowa State DNA Facility and PacBio sequencing was performed by the DNA Sequencing Center of Brigham Young University. Financial support for this research was provided by National Science Foundation award 1652661 (to D.G.S.).

## References

1. Grissa, I., Vergnaud, G. & Pourcel, C. The CRISPRdb database and tools to display CRISPRs and to generate dictionaries of spacers and repeats. BMC Bioinformatics 8, 172 (2007).

2. Godde, J. S. & Bickerton, A. The repetitive DNA elements called CRISPRs and their associated genes: evidence of horizontal transfer among prokaryotes. J Mol Evol 62, 718–729 (2006).

3. Makarova, K. S. et al. Evolutionary classification of CRISPR–Cas systems: a burst of class 2 and derived variants. Nature Reviews Microbiology 18, 67–83 (2020).

4. Sorek, R., Lawrence, C. M. & Wiedenheft, B. CRISPR-Mediated Adaptive Immune Systems in Bacteria and Archaea. Annual Review of Biochemistry 82, 237–266 (2013).

5. Mohanraju, P. et al. Diverse evolutionary roots and mechanistic variations of the CRISPR-Cas systems. Science 353, (2016).

6. Johnke, J. et al. Multiple micro-predators controlling bacterial communities in the environment. Curr Opin Biotechnol 27, 185–190 (2014).

7. Fuhrman, J. A. Marine viruses and their biogeochemical and ecological effects. Nature 399, 541–548 (1999).

8. Wommack, K. E., Ravel, J., Hill, R. T. & Colwell, R. R. Hybridization Analysis of Chesapeake Bay Virioplankton. Appl Environ Microbiol 65, 241–250 (1999).

9. Avrani, S., Wurtzel, O., Sharon, I., Sorek, R. & Lindell, D. Genomic island variability facilitates Prochlorococcus–virus coexistence. Nature 474, 604–608 (2011).

10. Deveau, H. et al. Phage Response to CRISPR-Encoded Resistance in Streptococcus thermophilus. Journal of Bacteriology 190, 1390–1400 (2008).

11. Datsenko, K. A. et al. Molecular memory of prior infections activates the CRISPR/Cas adaptive bacterial immunity system. Nat Commun 3, 945 (2012).

12. Sun, C. L. et al. Phage mutations in response to CRISPR diversification in a bacterial population. Environmental Microbiology 15, 463–470 (2013).

13. Rossetti, G. et al. The structural impact of DNA mismatches. Nucleic Acids Res 43, 4309–4321 (2015).

14. Sugimoto, N., Nakano, M. & Nakano, S. Thermodynamics−Structure Relationship of Single Mismatches in RNA/DNA Duplexes. Biochemistry 39, 11270–11281 (2000).

15. Borer, P. N., Dengler, B., Tinoco, I. & Uhlenbeck, O. C. Stability of ribonucleic acid double-stranded helices. Journal of Molecular Biology 86, 843–853 (1974).

16. Feng, H., Guo, J., Wang, T., Zhang, C. & Xing, X. Guide-target mismatch effects on dCas9–sgRNA binding activity in living bacterial cells. Nucleic Acids Research 49, 1263–1277 (2021).

17. Meeske, A. J., Nakandakari-Higa, S. & Marraffini, L. A. Cas13-induced cellular dormancy prevents the rise of CRISPR-resistant bacteriophage. Nature 1 (2019) doi:10.1038/s41586-019-1257-5.

18. Zheng, Y. et al. Endogenous Type I CRISPR-Cas: From Foreign DNA Defense to Prokaryotic Engineering. Frontiers in Bioengineering and Biotechnology 8, (2020).

19. Chylinski, K., Makarova, K. S., Charpentier, E. & Koonin, E. V. Classification and evolution of type II CRISPR-Cas systems. Nucleic Acids Res 42, 6091–6105 (2014).

20. Li, Y. et al. Cmr1 enables efficient RNA and DNA interference of a III-B CRISPR-Cas system by binding to target RNA and crRNA. Nucleic Acids Res 45, 11305–11314 (2017).

21. Wang, R. & Li, H. The mysterious RAMP proteins and their roles in small RNA-based immunity. Protein Sci 21, 463–470 (2012).

22. Kolesnik, M. V., Fedorova, I., Karneyeva, K. A., Artamonova, D. N. & Severinov, K. V. Type III CRISPR-Cas Systems: Deciphering the Most Complex Prokaryotic Immune System. Biochemistry Moscow 86, 1301–1314 (2021).

23. Vercoe, R. B. et al. Cytotoxic Chromosomal Targeting by CRISPR/Cas Systems Can Reshape Bacterial Genomes and Expel or Remodel Pathogenicity Islands. PLOS Genetics 9, e1003454 (2013).

24. Mojica, F. J. M., Díez-Villaseñor, C., García-Martínez, J. & Almendros, C. Short motif sequences determine the targets of the prokaryotic CRISPR defence system. Microbiology, 155, 733–740 (2009).

25. Sternberg, S. H., Redding, S., Jinek, M., Greene, E. C. & Doudna, J. A. DNA interrogation by the CRISPR RNA-guided endonuclease Cas9. Nature 507, 62–67 (2014).

26. Zetsche, B. et al. Cpf1 Is a Single RNA-Guided Endonuclease of a Class 2 CRISPR-Cas System. Cell 163, 759–771 (2015).

27. Horvath, P. et al. Diversity, Activity, and Evolution of CRISPR Loci in Streptococcus thermophilus. Journal of Bacteriology 190, 1401–1412 (2008).

28. Pattanayak, V. et al. High-throughput profiling of off-target DNA cleavage reveals RNA-programmed Cas9 nuclease specificity. Nature Biotechnology 31, 839–843 (2013).

29. Cong, L. et al. Multiplex Genome Engineering Using CRISPR/Cas Systems. Science 339, 819–823 (2013).

30. Künne, T., Swarts, D. C. & Brouns, S. J. J. Planting the seed: target recognition of short guide RNAs. Trends in Microbiology 22, 74–83 (2014).

31. Semenova, E. et al. Interference by clustered regularly interspaced short palindromic repeat (CRISPR) RNA is governed by a seed sequence. PNAS 108, 10098–10103 (2011).

32. Jiang, W., Bikard, D., Cox, D., Zhang, F. & Marraffini, L. A. RNA-guided editing of bacterial genomes using CRISPR-Cas systems. Nat Biotechnol 31, 233–239 (2013).

33. Maier, L.-K. et al. Essential requirements for the detection and degradation of invaders by the Haloferax volcanii CRISPR/Cas system I-B. RNA Biology 10, 865–874 (2013).

34. Wiedenheft, B. et al. RNA-guided complex from a bacterial immune system enhances target recognition through seed sequence interactions. PNAS 108, 10092–10097 (2011).

35. Tao, P., Wu, X. & Rao, V. Unexpected evolutionary benefit to phages imparted by bacterial CRISPR-Cas9. Science Advances 4, eaar4134 (2018).

36. Chabas, H. et al. Variability in the durability of CRISPR-Cas immunity. Philosophical Transactions of the Royal Society B: Biological Sciences 374, 20180097 (2019).

37. Pereira-Gómez, M. & Sanjuán, R. Effect of mismatch repair on the mutation rate of bacteriophage ϕX174. Virus Evolution 1, vev010 (2015).

38. Wu, X., Zhu, J., Tao, P. & Rao, V. B. Bacteriophage T4 Escapes CRISPR Attack by Minihomology Recombination and Repair. mBio 12, e01361–21.

39. Hossain, A. A., McGinn, J., Meeske, A. J., Modell, J. W. & Marraffini, L. A. Viral recombination systems limit CRISPR-Cas targeting through the generation of escape mutations. Cell Host & Microbe 29, 1482–1495.e12 (2021).

40. Mojica, F. J. M., Díez-Villaseñor, C., García-Martínez, J. & Soria, E. Intervening Sequences of Regularly Spaced Prokaryotic Repeats Derive from Foreign Genetic Elements. J Mol Evol 60, 174–182 (2005).

41. Bolotin, A., Quinquis, B., Sorokin, A. & Ehrlich, S. D. Clustered regularly interspaced short palindrome repeats (CRISPRs) have spacers of extrachromosomal origin. Microbiology 151, 2551–2561 (2005).

42. Heidelberg, J. F., Nelson, W. C., Schoenfeld, T. & Bhaya, D. Germ Warfare in a Microbial Mat Community: CRISPRs Provide Insights into the Co-Evolution of Host and Viral Genomes. PLOS ONE 4, e4169 (2009).

43. Semenova, E., Nagornykh, M., Pyatnitskiy, M., Artamonova, I. I. & Severinov, K. Analysis of CRISPR system function in plant pathogen Xanthomonas oryzae. FEMS Microbiol Lett 296, 110–116 (2009).

44. Pourcel, C., Salvignol, G. & Vergnaud, G. CRISPR elements in Yersinia pestis acquire new repeats by preferential uptake of bacteriophage DNA, and provide additional tools for evolutionary studies. Microbiology 151, 653–663 (2005).

45. Andersson, A. F. & Banfield, J. F. Virus Population Dynamics and Acquired Virus Resistance in Natural Microbial Communities. Science 320, 1047–1050 (2008).

46. Sun, C. L., Thomas, B. C., Barrangou, R. & Banfield, J. F. Metagenomic reconstructions of bacterial CRISPR loci constrain population histories. ISME J 10, 858–870 (2016).

47. Nussenzweig, P. M., McGinn, J. & Marraffini, L. A. Cas9 Cleavage of Viral Genomes Primes the Acquisition of New Immunological Memories. Cell Host & Microbe 26, 515–526.e6 (2019).

48. Fineran, P. C. et al. Degenerate target sites mediate rapid primed CRISPR adaptation. Proc Natl Acad Sci U S A 111, E1629–1638 (2014).

49. Xue, C. et al. CRISPR interference and priming varies with individual spacer sequences. Nucleic Acids Res 43, 10831–10847 (2015).

50. Xue, C., Whitis, N. R. & Sashital, D. G. Conformational Control of Cascade Interference and Priming Activities in CRISPR Immunity. Mol Cell 64, 826–834 (2016).

51. Cho, S. W. et al. Analysis of off-target effects of CRISPR/Cas-derived RNA-guided endonucleases and nickases. Genome Res. 24, 132–141 (2014).

52. Hsu, P. D. et al. DNA targeting specificity of RNA-guided Cas9 nucleases. Nature Biotechnology 31, 827–832 (2013).

53. Murugan, K., Seetharam, A. S., Severin, A. J. & Sashital, D. G. CRISPR-Cas12a has widespread off-target and dsDNA-nicking effects. J. Biol. Chem. jbc.R A120.012933 (2020) doi:10.1074/jbc.RA120.012933.

54. Strohkendl, I., Saifuddin, F. A., Rybarski, J. R., Finkelstein, I. J. & Russell, R. Kinetic Basis for DNA Target Specificity of CRISPR-Cas12a. Molecular Cell 71, 816–824.e3 (2018).

55. Jinek, M. et al. A Programmable Dual-RNA–Guided DNA Endonuclease in Adaptive Bacterial Immunity. Science 337, 816–821 (2012).

56. Jones, S. K. et al. Massively parallel kinetic profiling of natural and engineered CRISPR nucleases. Nature Biotechnology 1–10 (2020) doi:10.1038/s41587-020-0646-5.

57. Kleinstiver, B. P. et al. Genome-wide specificities of CRISPR-Cas Cpf1 nucleases in human cells. Nature Biotechnology 34, 869–874 (2016).

58. Endo, A., Masafumi, M., Kaya, H. & Toki, S. Efficient targeted mutagenesis of rice and tobacco genomes using Cpf1 from Francisella novicida. Sci Rep 6, 38169 (2016).

59. Alok, A. et al. The Rise of the CRISPR/Cpf1 System for Efficient Genome Editing in Plants. Frontiers in Plant Science 11, (2020).

60. Kim, D. et al. Genome-wide analysis reveals specificities of Cpf1 endonucleases in human cells. Nature Biotechnology 34, 863–868 (2016).

61. Zhong, Z. et al. Plant Genome Editing Using FnCpf1 and LbCpf1 Nucleases at Redefined and Altered PAM Sites. Mol Plant 11, 999–1002 (2018).

62. Chen, J. S. et al. CRISPR-Cas12a target binding unleashes indiscriminate single-stranded DNase activity. Science 360, 436–439 (2018).

63. Phan, P. T., Schelling, M., Xue, C. & Sashital, D. G. Fluorescence-based methods for measuring target interference by CRISPR-Cas systems. Methods Enzymol 616, 61–85 (2019).

64. Liu, Y. et al. Covalent Modifications of the Bacteriophage Genome Confer a Degree of Resistance to Bacterial CRISPR Systems. J Virol 94, e01630–20 (2020).

65. Barrangou, R. et al. CRISPR Provides Acquired Resistance Against Viruses in Prokaryotes. Science 315, 1709–1712 (2007).

66. van Houte, S. et al. The diversity-generating benefits of a prokaryotic adaptive immune system. Nature 532, 385–388 (2016).

67. Held, N. L. & Whitaker, R. J. Viral biogeography revealed by signatures in Sulfolobus islandicus genomes. Environ Microbiol 11, 457–466 (2009).

68. Zoephel, J. RNA-Seq analyses reveal CRISPR RNA processing and regulation patterns. Biochem Soc Trans 41, (2013).

69. Murugan, K., Suresh, S. K., Seetharam, A. S., Severin, A. J. & Sashital, D. G. Systematic in vitro specificity profiling reveals nicking defects in natural and engineered CRISPR–Cas9 variants. Nucleic Acids Research 49, 4037–4053 (2021).

70. Pyenson, N. C. & Marraffini, L. A. Co-evolution within structured bacterial communities results in multiple expansion of CRISPR loci and enhanced immunity. eLife 9, e53078 (2020).

71. Baba, T. et al. Construction of Escherichia coli K-12 in-frame, single-gene knockout mutants: the Keio collection. Mol Syst Biol 2, 2006.0008 (2006).

72. Jacob, F. & Wollman, E. L. [Processes of conjugation and recombination in Escherichia coli. I. Induction by conjugation or zygotic induction]. Ann Inst Pasteur (Paris) 91, 486–510 (1956).

73. Creating Excel files with Python and XlsxWriter — XlsxWriter Documentation. https://xlsxwriter.readthedocs.io/index.html.

