## Supplementary Information for "CRISPR-Cas effector specificity and target mismatches determine phage escape outcomes"

**Supplementary Figures**

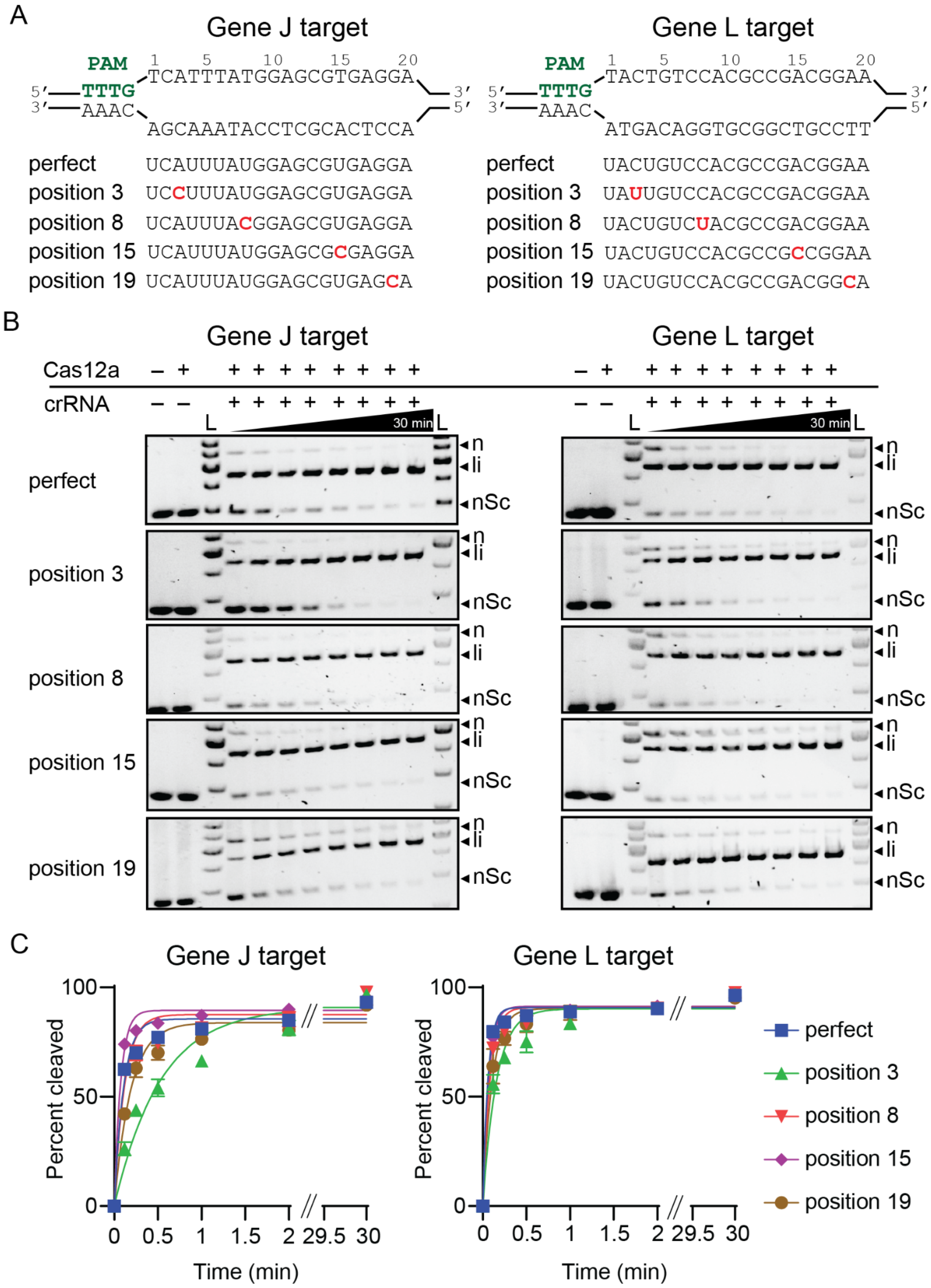

**Supplementary figure 1- Cleavage assays by FnCas12a with single mismatch crRNAs**

(A) Sequence of the target DNAs, perfectly matching crRNAs and single-mismatched crRNAs.
(B) Representative agarose gels showing time course cleavage of negatively supercoiled plasmid (nSC) using perfectly matching crRNA or single-mismatched crRNA by FnCas12a, resulting in linear (li) and/or nicked (n) products. Time points at which the samples were collected were 7 s, 15 s, 30 s, 1 min, 2 min, 5 min, 15 min, and 30 min. All controls were performed under the same conditions as the longest time point for the experimental samples.
(C) Quantification of cleaved products (linear and nicked fractions) from the time course cleavage. Averages of the cleaved fraction values were plotted versus time and fit to a first-order rate equation with error bars; n = 3 replicates. For values reported in Figure 1c, each individual replicate was fit, and $k_{obs}$ was reported as the average value for the three replicates.

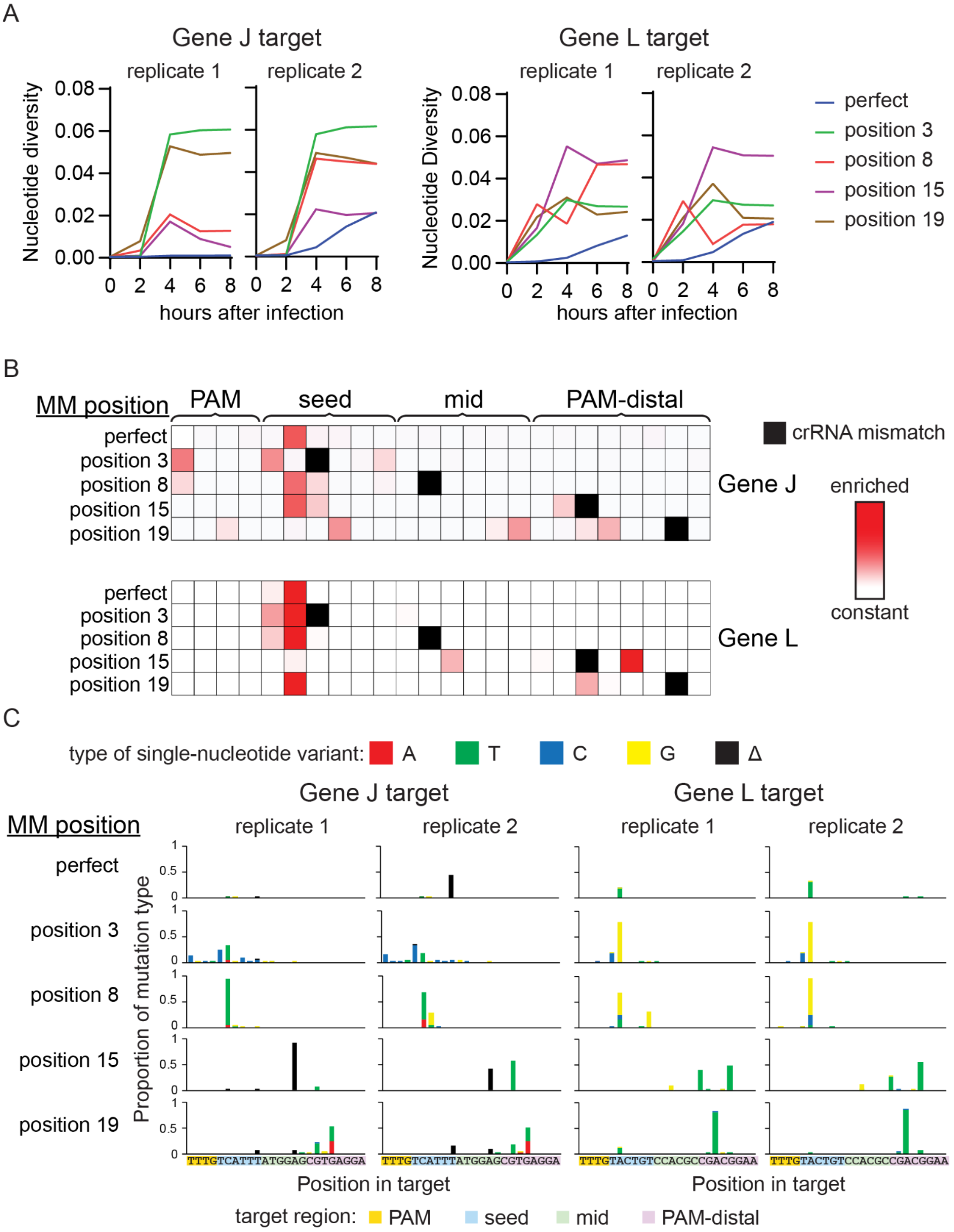

A
Gene J target
replicate 1
replicate 2
Nucleotide diversity
0.08
0.06
0.04
0.02
0.00
0 2 4 6 8
hours after infection
Gene L target
replicate 1
replicate 2
Nucleotide Diversity
hours after infection
perfect
position 3
position 8
position 15
position 19
B
MM position
PAM
seed
mid
PAM-distal
crRNA mismatch
perfect
position 3
position 8
position 15
position 19
Gene J
Gene L
enriched
constant
C
type of single-nucleotide variant: A T C G Δ
Gene J target
Gene L target
MM position
replicate 1
replicate 2
perfect
position 3
position 8
position 15
position 19
Proportion of mutation type
TTTGTCATTTATGGAGCGTGAGGA
TTTGTACTGTCCACGCCGACGGAA
Position in target
target region: PAM seed mid PAM-distal

**Supplementary figure 2 - Analysis of phage mutations that emerge following exposure to Cas12a-mediated interference with mismatched crRNAs**

(A) Line graphs showing the nucleotide diversity of phage target regions over time after exposure to bacteria cells expressing crRNAs with and without mismatches. Target regions are gene J or gene L and crRNAs either match the target region (perfect) or contain mismatches at position x. Nucleotide diversity is calculated using the proportion of each sequence in the sample and the number of nucleotide differences between each pair of sequences. Two individual replicates are shown for each condition.

(B) Heat maps showing location of target mutations that arose due to CRISPR targeting by FnCas12a on a solid medium. Z-scores for abundance of single-nucleotide variants, including nucleotide identity changes or deletions, were determined for each sample relative to the non-targeted control phage population. Enriched sequences indicate high Z-scores. Z-scores range from 0 (white) to 7.69 (darkest red). Single nucleotide deletions are shown at adjacent position to the 3' side. Positions with crRNA mismatches are labeled with solid black boxes in the heat map.

(C) Graphs showing single-nucleotide variations for mutated phage target sequences present at the 8 h time point for two individual replicates. Bar graph height shows the proportion of sequences in each sample with the mutation type at each position in the target. Deletions (Δ) are plotted at the first position where a mismatch occurs between the crRNA and the target.

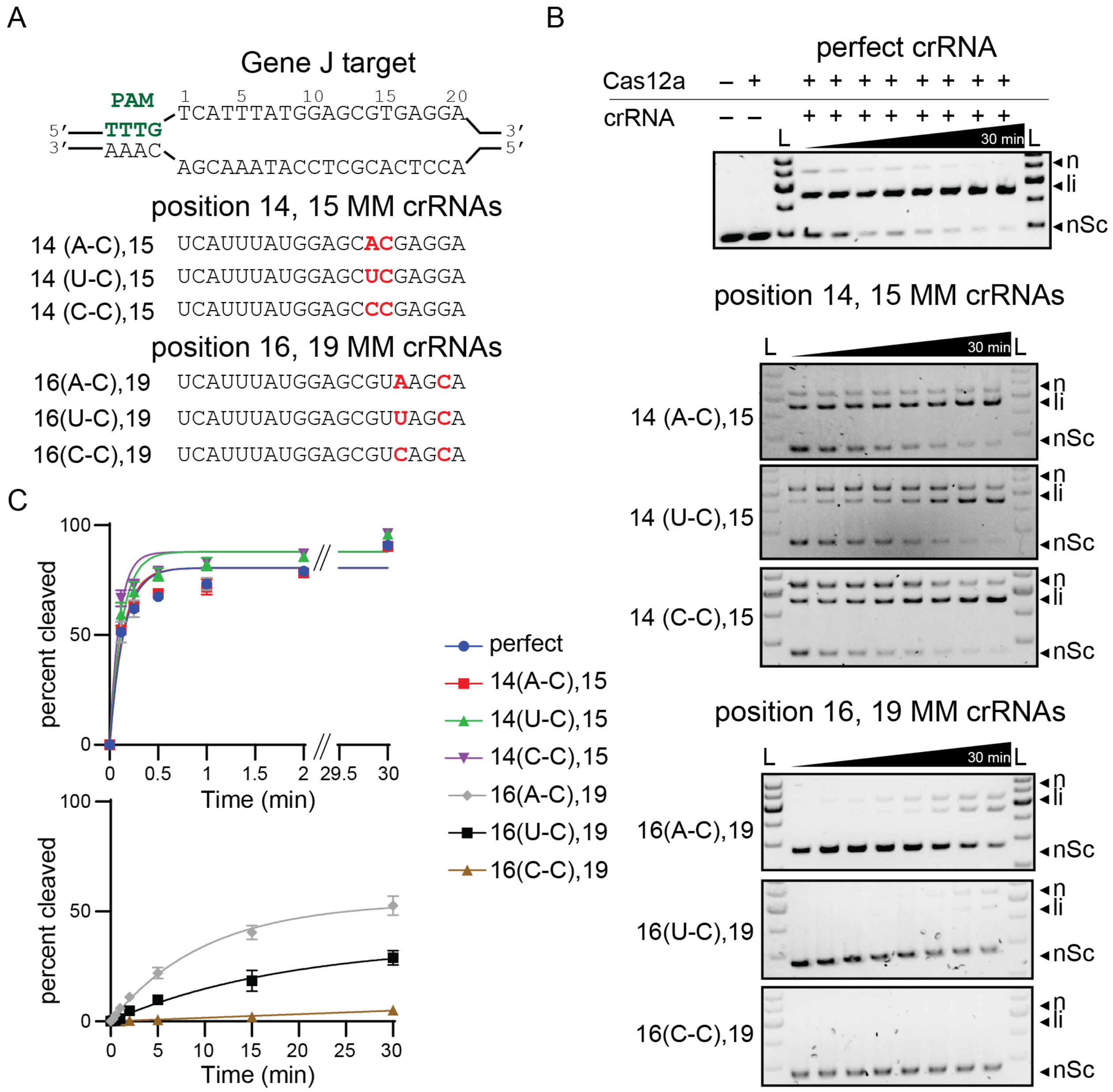


**Supplementary figure 3 - Cleavage assays by FnCas12a with double mismatch crRNAs**

(A) Sequence of the gene J target DNA, perfectly matching crRNA and double-mismatched crRNAs.

(B) Representative agarose gels showing time course cleavage of negatively supercoiled plasmid (nSC) using perfectly matching crRNA or double-mismatched crRNA by FnCas12a, resulting in linear (li) and/or nicked (n) products. Time points at which the samples were collected were 7 s, 15 s, 30 s, 1 min, 2 min, 5 min, 15 min, and 30 min. All controls were performed under the same conditions as the longest time point for the experimental samples.

(C) Quantification of cleaved products from the time course cleavage. The average cleaved fraction was plotted versus time and fit to a first-order rate equation with error bars representing standard deviation; n = 3 replicates.  For values reported in Figure 3b, each individual replicate was fit, and $k_{obs}$ was reported as the average value for the three replicates.

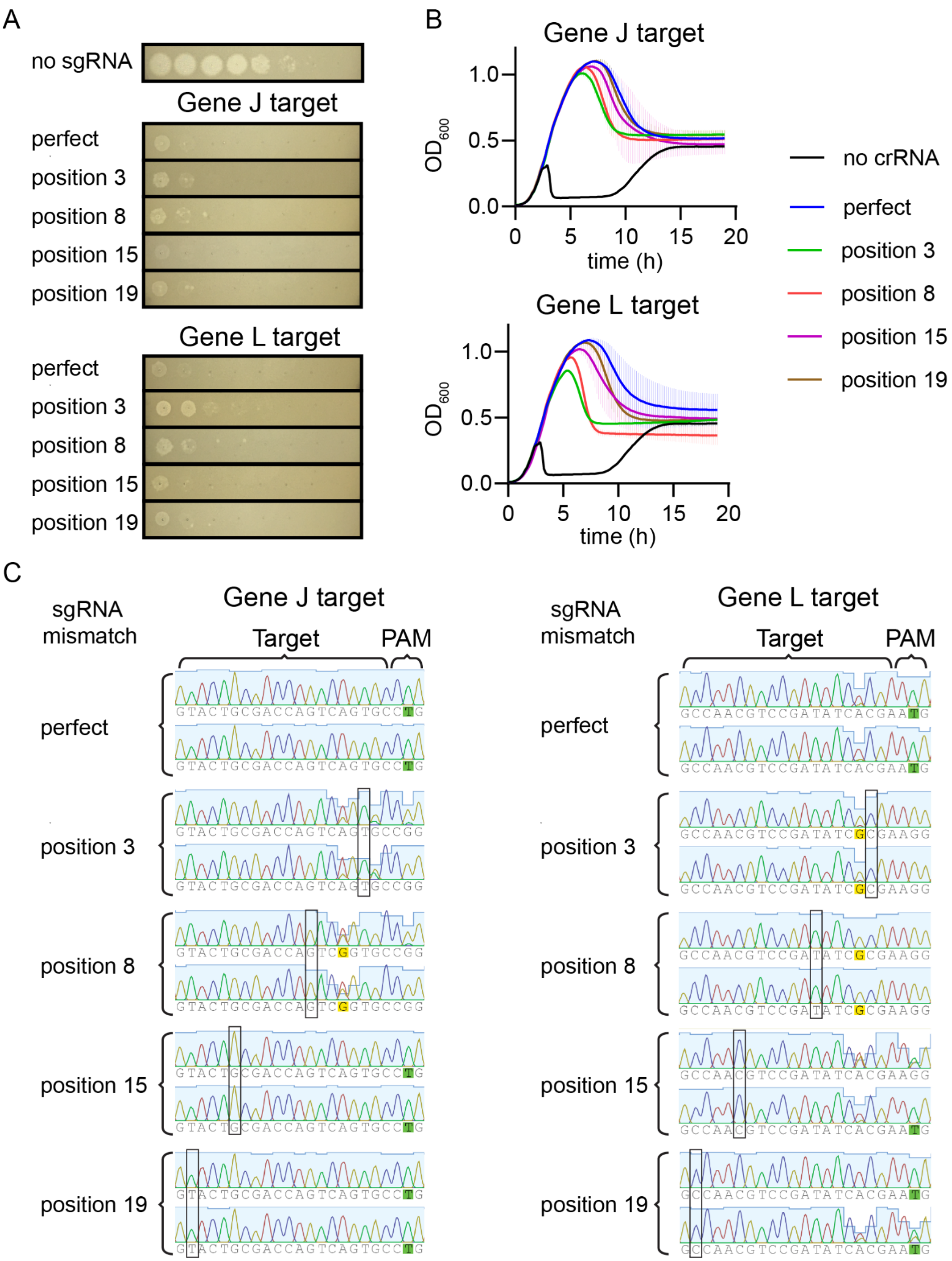
A
no sgRNA
Gene J target
perfect
position 3
position 8
position 15
position 19
Gene L target
perfect
position 3
position 8
position 15
position 19
B
Gene J target
Gene L target
OD600
1.0
0.5
0.0
0
5
10
15
20
time (h)
no crRNA
perfect
position 3
position 8
position 15
position 19
C
sgRNA mismatch
Gene J target
Target
PAM
perfect
GTACTGCGACCAGTCAGTGCCTG
GTACTGCGACCAGTCAGTGCCTG
position 3
GTACTGCGACCAGTCAGTGCCGG
GTACTGCGACCAGTCAGTGCCGG
position 8
GTACTGCGACCAGTCGGTGCCGG
GTACTGCGACCAGTCGGTGCCGG
position 15
GTACTGCGACCAGTCAGTGCCTG
GTACTGCGACCAGTCAGTGCCTG
position 19
GTACTGCGACCAGTCAGTGCCTG
GTACTGCGACCAGTCAGTGCCTG
sgRNA mismatch
Gene L target
Target
PAM
perfect
GCCAACGTCCGATATCACGAATG
GCCAACGTCCGATATCACGAATG
position 3
GCCAACGTCCGATATCGCGAAGG
GCCAACGTCCGATATCGCGAAGG
position 8
GCCAACGTCCGATATCGCGAAGG
GCCAACGTCCGATATCGCGAAGG
position 15
GCCAACGTCCGATATCACGAAGG
GCCAACGTCCGATATCACGAATG
position 19
GCCAACGTCCGATATCACGAATG
GCCAACGTCCGATATCACGAATG

**Supplementary figure 4 - Phage protection by SpCas9 with sgRNA mismatches**

(A) Spot assays using lambda phage on lawns of bacteria expressing SpCas9 along with sgRNAs with and without mismatches. SgRNAs target gene J or gene L and contain mismatches at position X or match the target (perfect).

(B) Growth curves of the same bacterial strains as in (A) after infection with lambda phage in liquid culture. Phage was added when the cells reached mid log phase at ~2 hours after inoculation. The average of three replicates is plotted for each condition, with error bars representing standard deviation.

(C) Sanger sequencing chromatograms of target regions (gene J or gene L) in phage populations targeted by SpCas9 armed with an sgRNA matching the target (perfect) or containing a mismatch at a position in the target. Position in the target is defined as the number of nucleotides away from the PAM going right to left, and is indicated with a box.

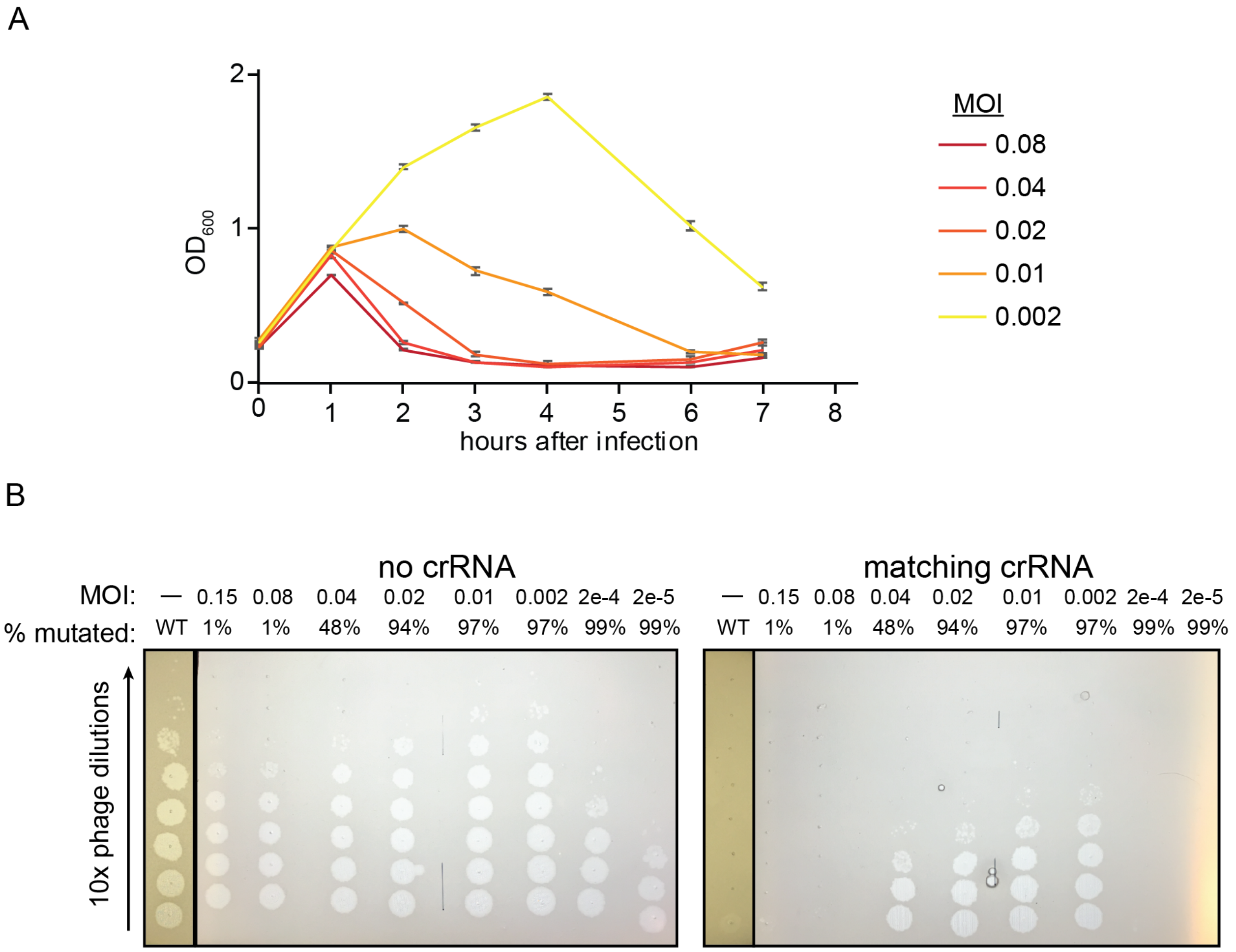


**Supplementary figure 5 - Mutant emergence at varied MOIs**

(A) Growth curves using cells expressing a crRNA targeting gene J with a mismatch in the seed region and infected with phage at different MOIs. Phage was added at the indicated MOIs when cells reached mid log phase and the $OD_{600}$ of the culture was measured over time. Cultures at lower MOIs did not lyse and are omitted from the graph.

(B) Spot assays estimating the titer of phage lysates exposed to interference by Cas12a armed with a seed mismatched crRNA. Full plates from Fig. 4b, including lowest MOI samples which produced phages with low titers.

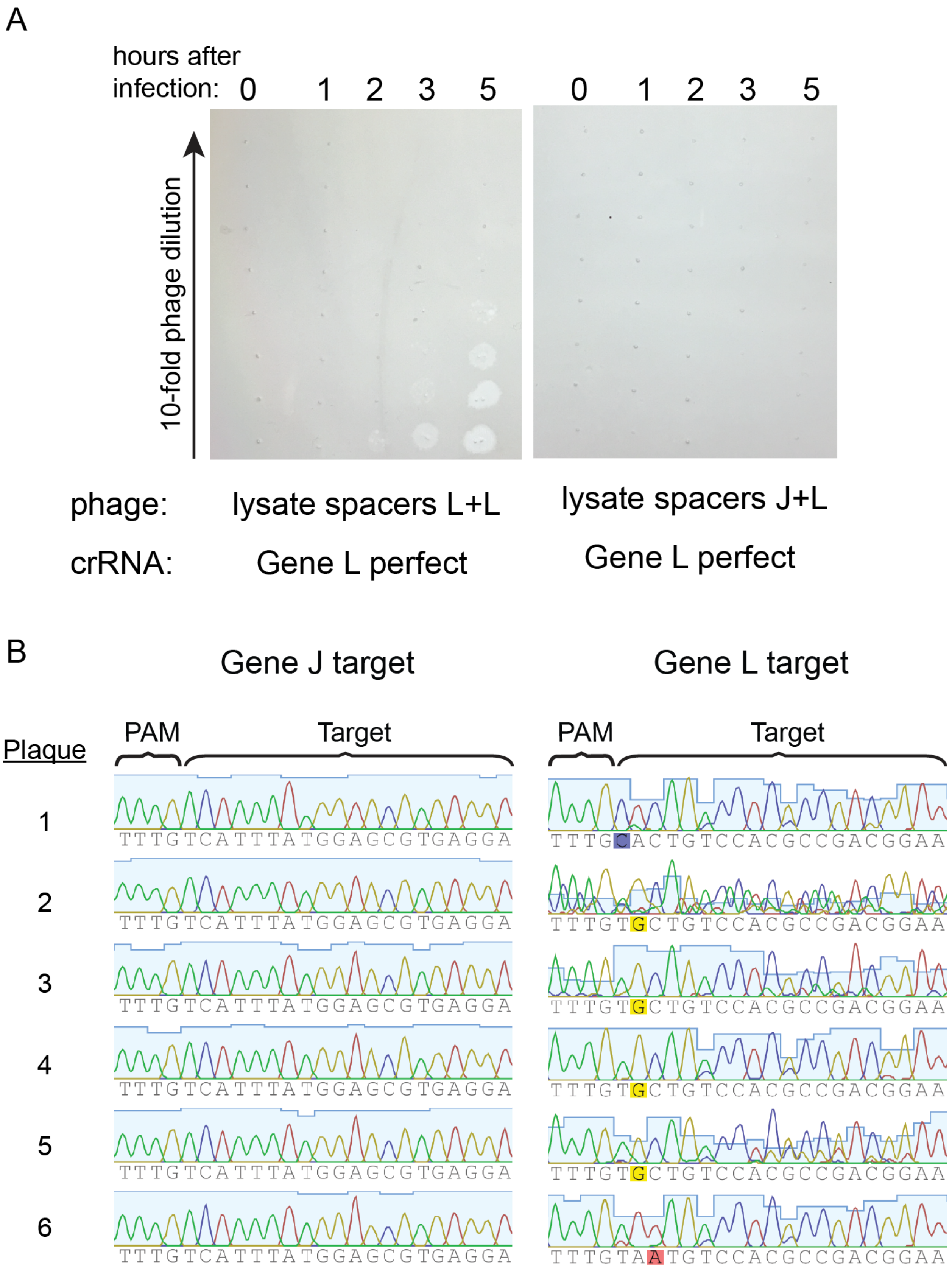
A
hours after infection: 0 1 2 3 5
0 1 2 3 5
10-fold phage dilution
phage: lysate spacers L+L
lysate spacers J+L
crRNA: Gene L perfect
Gene L perfect
B
Gene J target
Gene L target
Plaque
PAM
Target
1
2
3
4
5
6
TTTGTCATTTATGGAGCGTGAGGA
TTTGCACTGTCCACGCCGACGGAA
TTTGTGCTGTCCACGCCGACGGAA
TTTGTAATGTCCACGCCGACGGAA

**Supplementary figure 6 - Phage targeted by multiple spacers develop mutations in one or more targeted regions**

(A) Spot assays performed using lambda phage that previously infected *E. coli* in liquid culture expressing FnCas12a and two different crRNAs targeting gene J and L (lysate spacers J+L) or two of the same crRNA targeting gene L (lysate spacers L+L) both with mismatches in the seed region. Phage was harvested at different time points of the liquid culture (0,1,2,3 and 5 hours after infection). The previous phage lysates were spotted on cells expressing a crRNA that matches the gene L target in the lambda genome (gene L perfect)

(B) Sanger sequencing chromatograms showing sequences of target regions of the genome in phage exposed to cells expressing two mismatched crRNAs targeting gene J and gene L respectively. Phage from single plaques were isolated and both target regions were sequenced. Mutated bases are highlighted.

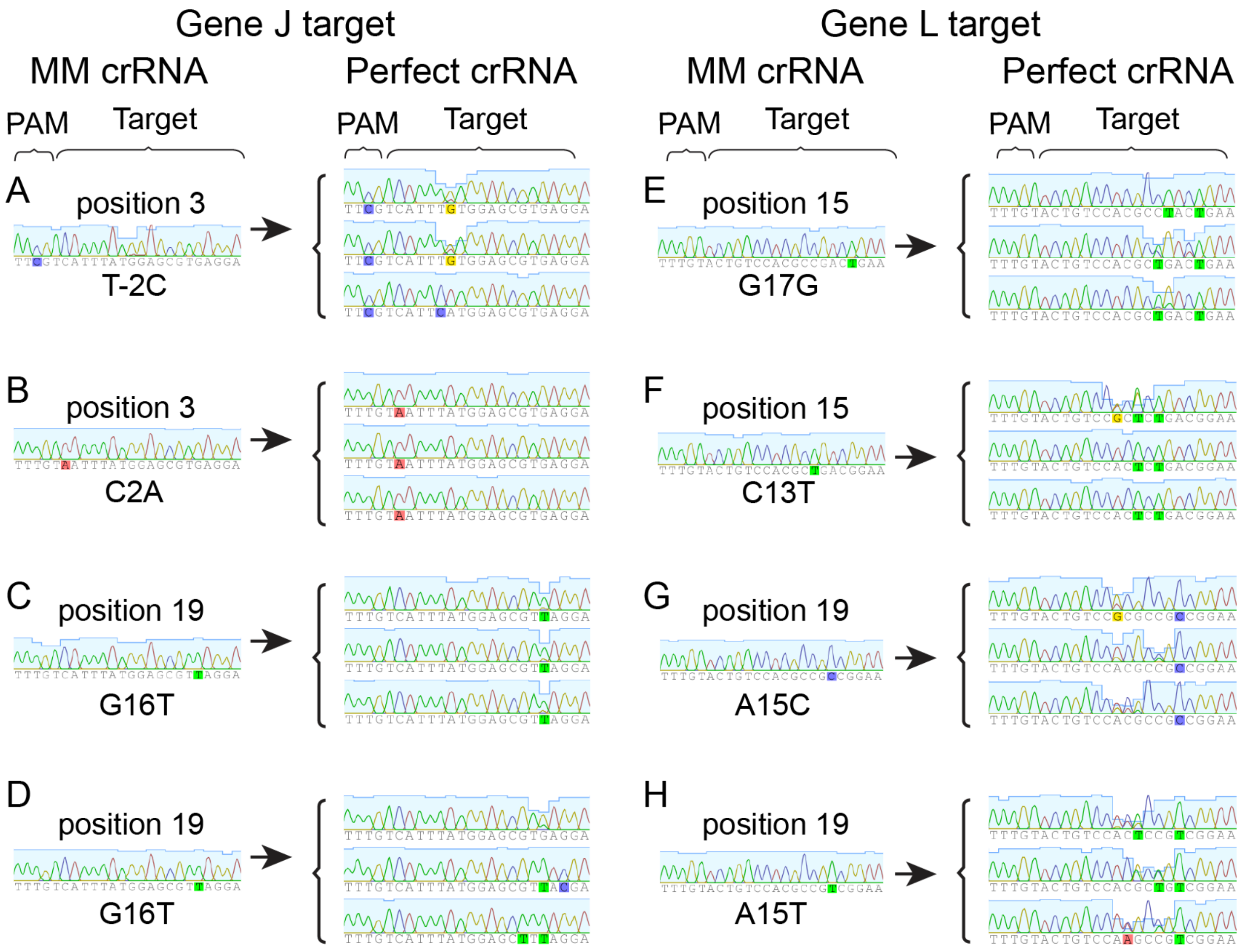


**Supplementary figure 7 - Purified single mutant and double mutant chromatograms**

(A-H) Sanger sequencing chromatograms of single and double mutant phage lysates. Single mutant phage were generated by exposure to crRNAs with mismatches (MM crRNA) at different positions (position X) and purified as shown in Fig. 6a. Mutants were generated in the gene J and gene L CRISPR target. Purified single mutant phage were used to challenge bacteria expressing a perfect crRNA and target regions were sequenced by Sanger sequencing to determine if second mutations appeared. Both mixed and clonal double mutant populations were generated after this step. Mutated bases are highlighted.

A

| crRNA | Gene J target number of reads rep 1 | Gene J target number of reads rep 2 | Gene L target number of reads rep 1 | Gene L target number of reads rep 2 |
|---|---|---|---|---|
| non-targeting | 644959 | 240080 | 702921 | 627352 |
| perfect | 40833 | 47381 | 35357 | 44965 |
| position 3 | 91218 | 86032 | 74030 | 68508 |
| position 8 | 83682 | 82296 | 70347 | 72941 |
| position 15 | 89176 | 85471 | 72068 | 66747 |
| position 19 | 90221 | 75516 | 88252 | 78551 |

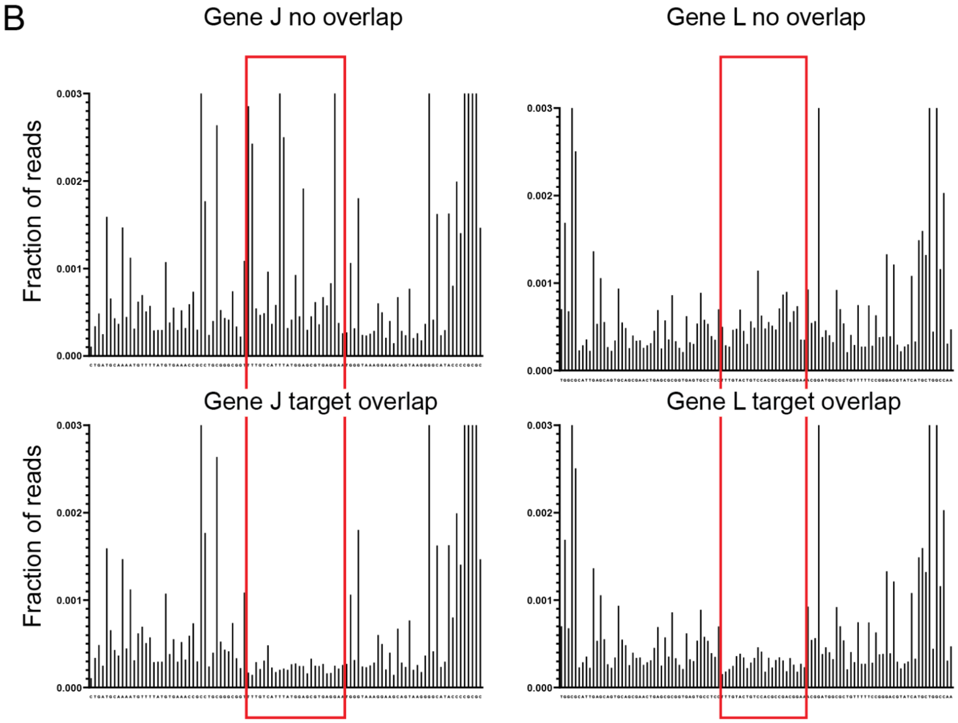


**Supplementary figure 8 - MiSeq sample counts and R1/R2 file overlap**

(A) Table showing absolute counts from MiSeq for each replicate of the 8 h time point for each experimental condition. Each count represents an extracted sequence in which R1 and R2 reads matched. The negative control (non-targeting crRNA) samples were run in a separate MiSeq run to maximize the number of reads and minimize barcode overlap with mutated samples, allowing for analysis of pre-existing mutants in the wild-type population.

(B) Bar charts showing mutated sequences at each position in the high-throughput sequencing reads of the negative control lambda phage population for the gene J and gene L region. The target region is highlighted with a red box. R1 and R2 reads do not overlap in the target region (no overlap) or overlap in the target region (target overlap). R1 reads are used for the target region in the no overlap condition. When R1 and R2 reads overlap, sequences in which the target region sequence does not agree for both the R1 and R2 reads are removed from analysis and are not shown in this figure. This measure was taken to ensure that variations observed in negative control samples arose solely from PCR errors or the natural variation of the population.

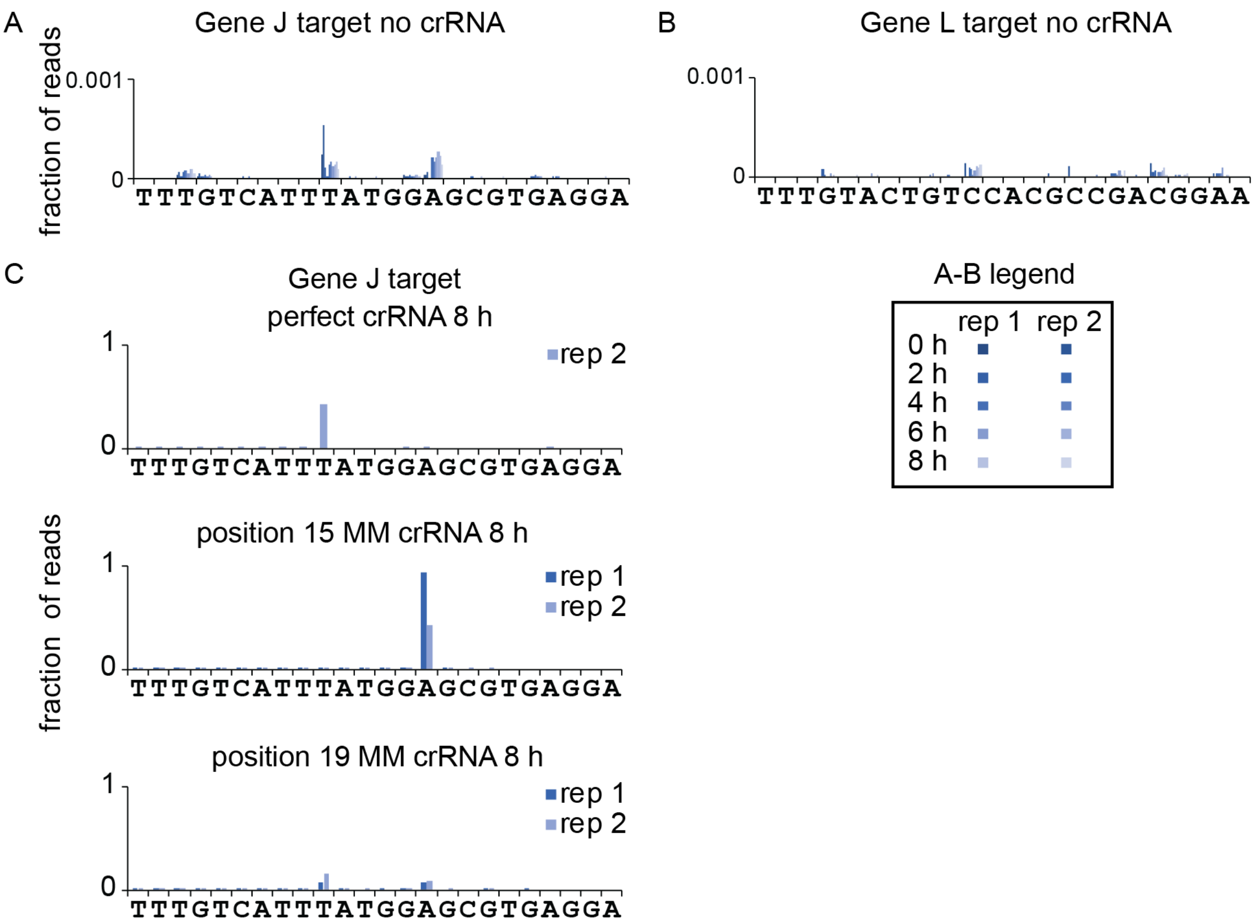


**Supplementary figure 9 - Single deletions enriched by CRISPR exposure**

Bar charts showing single nucleotide deletions from the lambda phage gene J target (A) and gene L target (B) in phage that were exposed to cells expressing a non-targeting crRNA (CRISPR inactive) and cells expressing crRNAs with and without mismatches to the lambda phage gene J target (C). (A-B) Deletions are mapped along the target sequences for all time points and both biological replicates for the negative control samples. (C) CRISPR active samples shown for gene J target that contained deletion sequences that represented more than 1% of all reads. No such deletions were observed in the gene L target. Deletions were observed in the gene L coding region in phage in the natural population. The deletions could remain in genomes in the population as these genomes are packaged along with functional structural proteins in successfully infected cells.

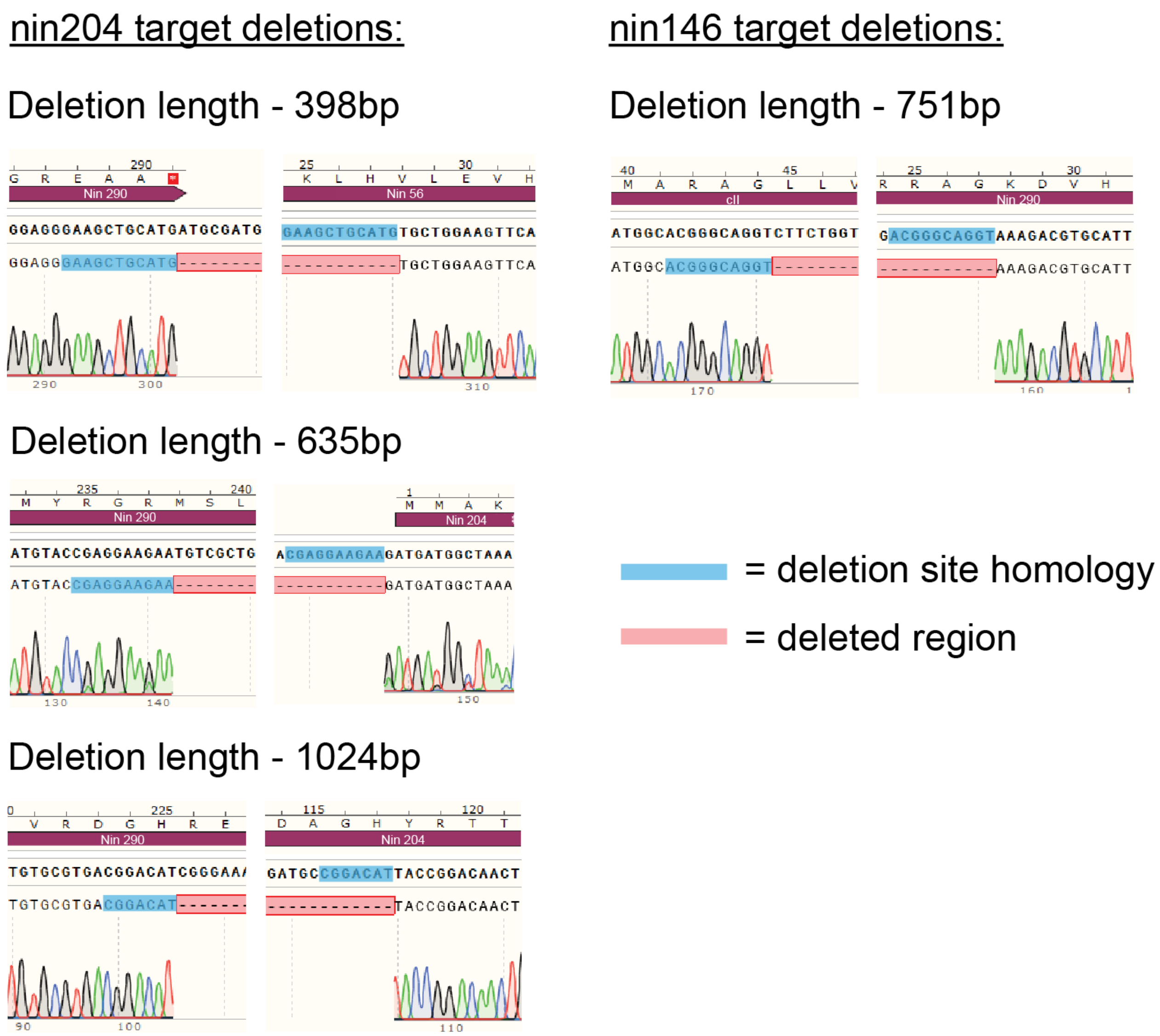


**Supplementary figure 10 - CRISPR escape deletions in non-essential regions**

Sanger sequencing chromatograms of phage genome deletions in non-essential regions targeted by Cas12a. Non-essential regions in the lambda phage genome were targeted with matching crRNAs on solid media. Phages were isolated and the target regions were PCR amplified. Single bands were gel purified and PCR amplified in a second round. These second PCR products were sequenced and the chromatograms were aligned to the WT lambda phage genome. Homology at the each end of the deletions was identified and highlighted in blue.
